# Whole-genome sequencing uncovers cryptic and hybrid species among Atlantic and Pacific cod-fish

**DOI:** 10.1101/034926

**Authors:** Katrín Halldórsdóttir, Einar Árnason

## Abstract

Speciation often involves the splitting of a lineage and the adaptation of daughter lineages to different environments. It may also involve the merging of divergent lineages, thus creating a stable homoploid hybrid species^1^ that constructs a new ecological niche by transgressing^2^ the ecology of the parental types. Hybrid speciation may also contribute to enigmatic and cryptic biodiversity in the sea.^3,4^ The enigmatic walleye pollock, which is not a pollock at all but an Atlantic cod that invaded the Pacific 3.8 Mya,^5^ differs considerably from its presumed closest relatives, the Pacific and Atlantic cod. Among the Atlantic cod, shallow-water coastal and deep-water migratory frontal ecotypes are associated with highly divergent genomic islands;^6,7^ however, intermediates remain an enigma.^8^ Here, we performed whole-genome sequencing of over 200 individuals using up to 33 million SNPs based on genotype likelihoods^9^ and showed that the evolutionary status of walleye pollock is a hybrid species: it is a hybrid between Arctic cod and Atlantic cod that transgresses the ecology of its parents. For the first time, we provide decisive evidence that the Atlantic cod coastal and frontal ecotypes are separate species that hybridized, leading to a true-breeding hybrid species that differs ecologically from its parents. We refute monophyly and dichotomous branching of these taxa, and stress the importance of looking beyond branching trees at admixture and hybridity. Our study demonstrates the power of whole-genome sequencing and population genomics in providing deep insights into fundamental processes of speciation. Our study was a starting point for further work aimed at examining the criteria of hybrid speciation,^10^ selection, sterility and structural chromosomal variation^11^ among cod-fish, which are among the most important fish stocks in the world. The hybrid nature of both the walleye pollock and Atlantic cod raises the question concerning the extent to which very profitable fisheries^12,13^ depend on hybrid vigour. Our results have implications for management of marine resources in times of rapid climate change.^14,15^

---

Speciation often involves the splitting of a lineage and the adaptation of daughter lineages to different environments. It may also involve the merging of divergent lineages, thus creating a stable homoploid hybrid species^1^ that constructs a new ecological niche by transgressing^2^ the ecology of the parental types. Hybrid speciation, which is well known among plants,^16,17^ is found also among animals, such as *Heliconius* butterflies^18^ and swordfish.^10,19^ Models of homoploid hybrid speciation involve chromosomal rearrangements and reduced fertility of *F_1_* hybrids, which may produce novel balanced gametes. Inbreeding of the *F_1_* may lead to *F*_2_ individuals with novel fertile and stable homokaryotypes that are, at least partially, reproductively isolated from the parental types.^16,17^ We suggest that hybrids among high-fecundity promiscuously mating organisms, such as cod-fish, could also pass through the *F*_1_ barrier. Ecological selection is also very important for the climbing of a new adaptive peak by hybrid species.^1,20^

Cryptic and sibling species, forms that are very similar morphologically, are common in the sea and may reflect adaptive divergence of habitat use, life-history and chemical recognition without morphological divergence.^3,21^ Marine populations often have high dispersal potential and the marine environment appears to have few barriers to gene flow. Thus allopatric divergence may be slow. Speciation in the marine environment frequently involves behavioural differences in spawning time and mate recognition, gametic incompatibility and habitat specialization such as salt tolerance.^22^ The role of hybrid speciation^1^ contributing to enigmatic and cryptic biodiversity in the sea^3,4^ is unknown. The origin of morphologically distinct forms and niche shifts may present an enigma. Morphologically similar forms are often cryptic species that are genetically distinct as revealed molecular genetic studies.^4^ Identifying cryptic species is important for evaluation of biodiversity. Identifying cryptic species complexes in commercially exploited organisms also is important for conservation and the protection and management of natural resources.

Cod-fish represent some of the most important commercial fisheries in the world. Among cod-fish, the enigmatic walleye pollock (*Gadus chalcogrammus*), which is not a pollock at all, is under the hypothesis of speciation by lineage-splitting of an Atlantic cod (*Gadus morhua*) that invaded the Pacific Ocean 3.8 Mya, according to mtDNA genomics.^5^ Pacific cod (*Gadus macrocephalus*) is a slightly older (4 Mya) invasion.^5^ However, Pacific and Atlantic cod share more traits than either of them share with walleye pollock. The semi-pelagic schooling walleye pollock differs morphologically, ecologically and behaviourally from these presumed closest relatives. The specific traits of walleye pollock niche shift would then have to have arisen by selective filtering during colonization or subsequent adaptation to Pacific environments. However, the Pacific cod that colonized the same habitat did not go through the same filtering. The Pacific cod is also thought to have re-invaded the Arctic and Atlantic Oceans at western Greenland and formed the Greenland cod *Gadus ogac.^5^* The Greenland cod is morphologically similar to both the Pacific and Atlantic cod and thus no special filtering occurred at re-invasion under that hypothesis. The biogeography of these taxa makes walleye pollock stand out as an evolutionary enigma.

Among Atlantic cod, the shallow-water coastal and deep-water frontal behavioural ecotypes, which are defined by storage-tag data,^23,24^ correlate with *AA* and *BB* homozygotes, respectively, of the *Pan* I locus^23,25^ located in a highly divergent genomic island on linkage group LG01.^6,7,26^ However, there is a general heterozygote excess at this locus.^27–29^ Heterozygotes and some homozygotes are behaviourally atypical, being intermediate, for both the coastal and frontal types.^23^ A study of other genomic islands suggests cryptic speciation and extensive hybridization producing *F*_1_ hybrids and few, if any, *F*_2_ and back-crossed individuals.^8^ Thus, if the coastal and frontal types are reproductively isolated, as has been suggested,^7,30,31^ the intermediates are an enigma: is most of the population composed of sterile hybrids, as implied by the lack of *F*_2_ and back-crossed individuals^8^? This is hardly a tenable proposition.

Population genomics promises to significantly advance our knowledge of enigmatic and cryptic forms and their speciation in the sea. Here, we performed whole-genome sequencing of over 200 individual cod-fish using up to 33 million SNPs based on genotype likelihoods^9^ to elucidate evolutionary relationships and speciation among gadid taxa.

## Results and Discussion

An individual admixture analysis differentiated the Atlantic, Pacific and Arctic taxa with a model of *k =* 2 ancestral populations (Figure 1). However, the walleye pollock genome of all individuals was about 40% Atlantic cod and 60% Arctic cod (*Boreogadus saida*). As the number of ancestral populations in the model increased, the groups split up: western vs eastern Atlantic (*k* = 3), Pacific and Greenland cod vs Arctic cod, walleye pollock and frontal ecotype vs coastal ecotype (*k* = 4) and the coastal ecotype into north vs south (*k* = 5). However, walleye pollock was always about 40% Atlantic cod, except at *k =* 4, where it aligns with Arctic cod. High-coverage data showed similar patterns (although the proportions differed; Figure 2) that the pollock genome was 50%, 10% and 40% Atlantic, Pacific and Arctic cod, respectively. Walleye pollock was similarly admixed at all genomic regions assignable to linkage groups (Supplemental Figure 1). As a side-note, a single individual of Polar cod (*Arctogadus glacialis*) was similarly admixed, possibly from similar processes as those detected in walleye pollock. The first and second principal components (PCs) separated Pacific and Greenland cod on one linear cluster, Atlantic cod on another and Arctic cod on a third (Figure 3). Greenland cod clusters with Pacific cod (Supplemental Figure 2) but in general it is closer to the other taxa by the analysis of principal components (Figure 3). The linear behaviour represented geographic variation within each species. The walleye pollock (and the single Polar cod individual) lay at the nexus of the other species.

**Figure 1.**
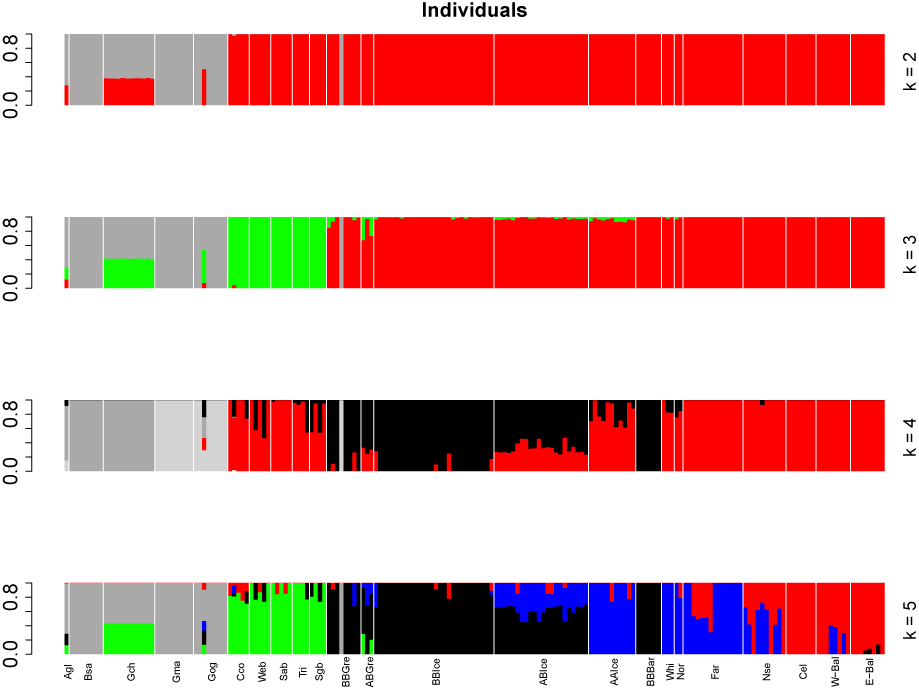
Individual admixture analysis of cod-fish. Ordinate is individual admixture proportion in a given model. The Number of ancestral populations *k* in the model range from 2 to 5. Results ordered based on species *Arctogadus glacialis* Agl, *Boreogadus saida* Bsa, *Gadus chalcogrammus* Gch, *Gadus macrocephalus* Gma, *Gadus ogac* Gog, and for *Gadus morhua* by localities from west to east and by ecotype: Cape cod Cco, Western Bank Web, Sable Bank Sab, Trinity Bay Tri, Southern Grand Banks Sgb, Greenland frontal BBGre, Greenland intermediate ABGre, Iceland frontal BBIce, Iceland intermediate ABIce, Iceland coastal AAIce, Barents Sea frontal BBBar, White Sea Whi, Norway Nor, Faroe Islands Far, North Sea Nse, Celtic Sea Cel, western Baltic W-Bal, and eastern Baltic E-Bal. Based on 15 million variable sites from the entire genome after filtering.

**Figure 2.**
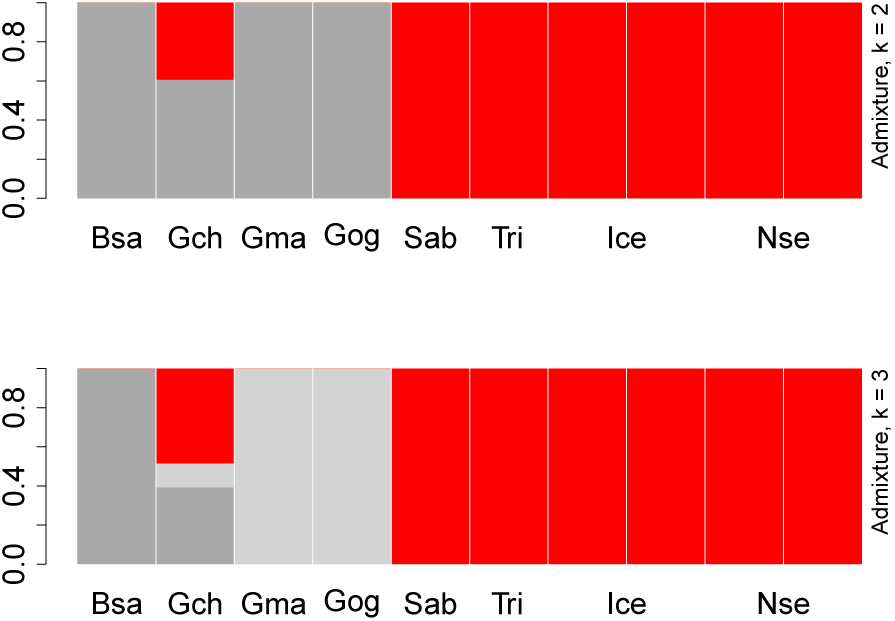
Individual admixture analysis of Pacific, Arctic and Atlantic cod-fish. Based on whole genome sequencing with 20 – 30 × coverage of entire genome. *Boreogadus saida* Bsa, *Gadus chalcogrammus* Gch, *Gadus macrocephalus* Gma, *Gadus ogac* Gog, and *Gadus morhua* Gmo individuals from Sable Bank Sab, Trinity Bay Tri, Iceland Ice, and North Sea Nse. Based on 32.9 million variable sites from the entire genome after filtering.

**Figure 3.**
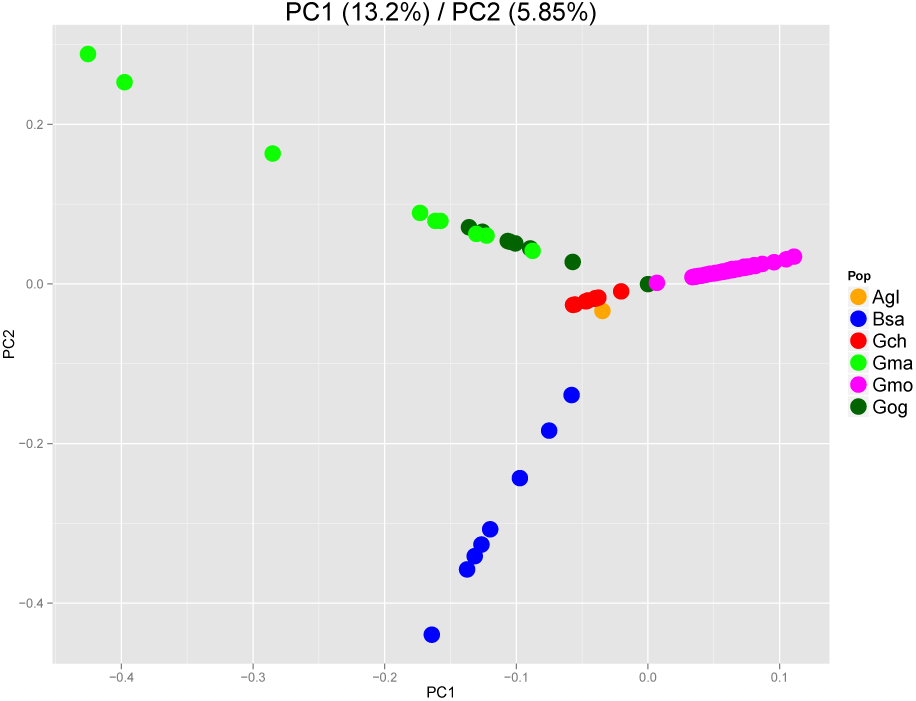
Principal component analysis (PCA) of genomic variation of six species of Atlantic, Arctic, and Pacific cod-fish. First and second principal components. *Arctogadus glacialis* Agl, *Boreogadus saida* Bsa, *Gadus chalcogrammus* Gch, *Gadus macrocephalus* Gma, *Gadus morhua* Gmo, *Gadus ogac* Gog.

Our interpretation of the evidence was that walleye pollock is a hybrid between Atlantic and Arctic cod. Walleye pollock shares two morphological traits with Arctic cod: an absent or much reduced chin barbel sensory organ and a forked tail, which define genera (sensu Svetovidov, 1948) within the Gadinae.^32^ Fisheries survey experts have difficulty in distinguishing between older slow-growing Arctic cod and younger fast-growing walleye pollock.^33^ These facts independently support our thesis that walleye pollock is a homoploid hybrid species.

The minimum evolutionary tree of whole-genome genetic distances (Supplemental Figure 2) had the same topology as the mtDNA tree.^5^ However, this result was dependent on monophyly of speciation by lineage splitting. We refuted this hypothesis in the case of walleye pollock. Furthermore, the timing^5^ of the colonization of the Pacific by walleye pollock cannot be estimated from the dichotomous tree. Our analysis implies that it might have happened even as recently as 200 years ago, when the species was discovered and described. We also question the biogeographical hypothesis that Pacific cod is of an Atlantic cod invasion of the Pacific ocean and that Greenland cod is of a re-invasion of the Atlantic by Pacific cod.^5^ A more parsimonious single-invasion biogeographical hypothesis is that Greenland cod is a speciation from Atlantic cod at Greenland and that Pacific cod is of an invasion of Greenland cod into the Pacific ocean (see Figure 1 and Supplemental Figure 2).

Among Atlantic cod, an individual admixture analysis revealed western cod, eastern cod and the frontal ecotype as separate entities (*k* = 3) (Figure 4 and Supplemental Figure 1). All three remain distinct entities at all *k* values, and there is a hybrid zone in the western Baltic.^34^ The coastal fish splits into Icelandic, White Sea, Norway and North Sea fish on the one hand, and the Faroe Islands and the Celtic Sea on the other, and the Baltic cod is clearly divergent (Supplemental Figure 3). The genomes of the intermediate (*Pan* I *AB*) of Greenland and Iceland are about 50% frontal and 50% coastal ecotypes (Figure 4, *k =* 5). However, the *Pan* I locus, which is a proxy for behavioural ecotypes,^23,24^ is located in a large genomic island of divergence on linkage group LG01.^6,7,26^ Thus, variation in LG01 may have overriding influence on the admixture patterns (Figure 4).

**Figure 4.**
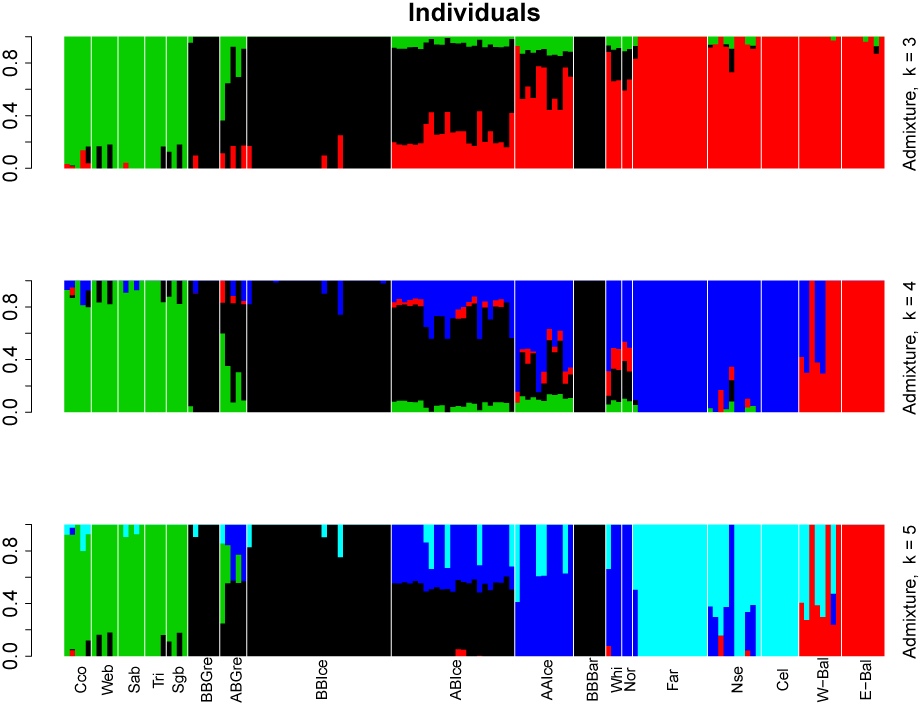
Admixture analysis of individual Atlantic cod. Number of ancestral populations *k* ranging from 3 to 5. Results ordered from west to east and by *Pan* I genotype proxy for behavioural ecotype: Cape cod Cco, Western Bank Web, Sable Bank Sab, Trinity Bay Tri, Southern Grand Banks Sgb, Greenland frontal BBGre, Greenland intermediate ABGre, Iceland frontal BBIce, Iceland intermediate ABIce, Iceland coastal AAIce, Barents Sea frontal BBBar, White Sea Whi, Norway Nor, Faroe Islands Far, North Sea Nse, Celtic Sea Cel, western Baltic W-Bal, and eastern Baltic E-Bal. Based 8.6 million variable sites from the entire genome.

To address this question, we analysed separately the LG02 to LG23 linkage groups, and then added LG01. We also analysed genomic regions that did not map to linkage groups (see methods).

The assessment of population differentiation by sliding-window *F_ST_* between north and south in the eastern Atlantic (Extended Data Figure 4) revealed extensive islands of divergence in LG01, LG02, LG07 and LG12, as observed previously.^6,8^ However, the LG01 and LG07 islands were complex, as they were divided by subregions showing no differentiation, thus indicating the independence of the subregions. Most other linkage groups also showed smaller regions (mini islands) of high differentiation. The comparison of north and south with west revealed a higher level of differentiation throughout the genome, in accordance with the admixture analysis, which suggests that western cod is a distinct entity. The LG07 island appeared in north/south and west/south but not in west/north comparisons. Therefore, it evolved in the south (coastal ecotype). Similarly, the LG01 island characterized north (frontal/intermediate) and the LG12 island differed between north and south. The fact that genomic islands differed between geographic regions has implications for the interpretation of long-distance linkage disequilibrium.^8^

There was considerable geographic variation. The west was clearly different from the east (Supplemental Figure 5), and the eastern Baltic stood out (Supplemental Figure 3), which is suggestive of species level differentiation of the western cod and of the eastern Baltic cod. From this evidence they are cryptic species within the Atlantic cod complex. To eliminate or reduce the effects of geographic variation, we focused on behavioural ecotypes from Iceland. A discriminant analysis of principal components (DAPC) using LG01 *Pan* I genotype priors showed the presence of three clusters for genomic regions LG02 to LG23, the part of the genome that definitely does not map to LG01 (Figure 5): a cluster of *AA* homozygotes only, a cluster of *BB* homozygotes only and a third and larger cluster of all *AB* heterozygotes, together with some *AA* and *BB* individuals. The addition of LG01 to the analysis retained the three groups with homozygotes of the heterogeneous group separating somewhat on the first but not on the second discriminant function. Similarly, the remaining genome that does not map to linkage groups showed the same pattern. In essence, there was an intra-class correlation within groups defined by linkage group LG01 in the parts of the genome that did not contain LG01, which implies the existence of a correlation between linkage groups (cf. long-distance linkage disequilibrium^8^) that also extends to the whole genome.

**Figure 5.**
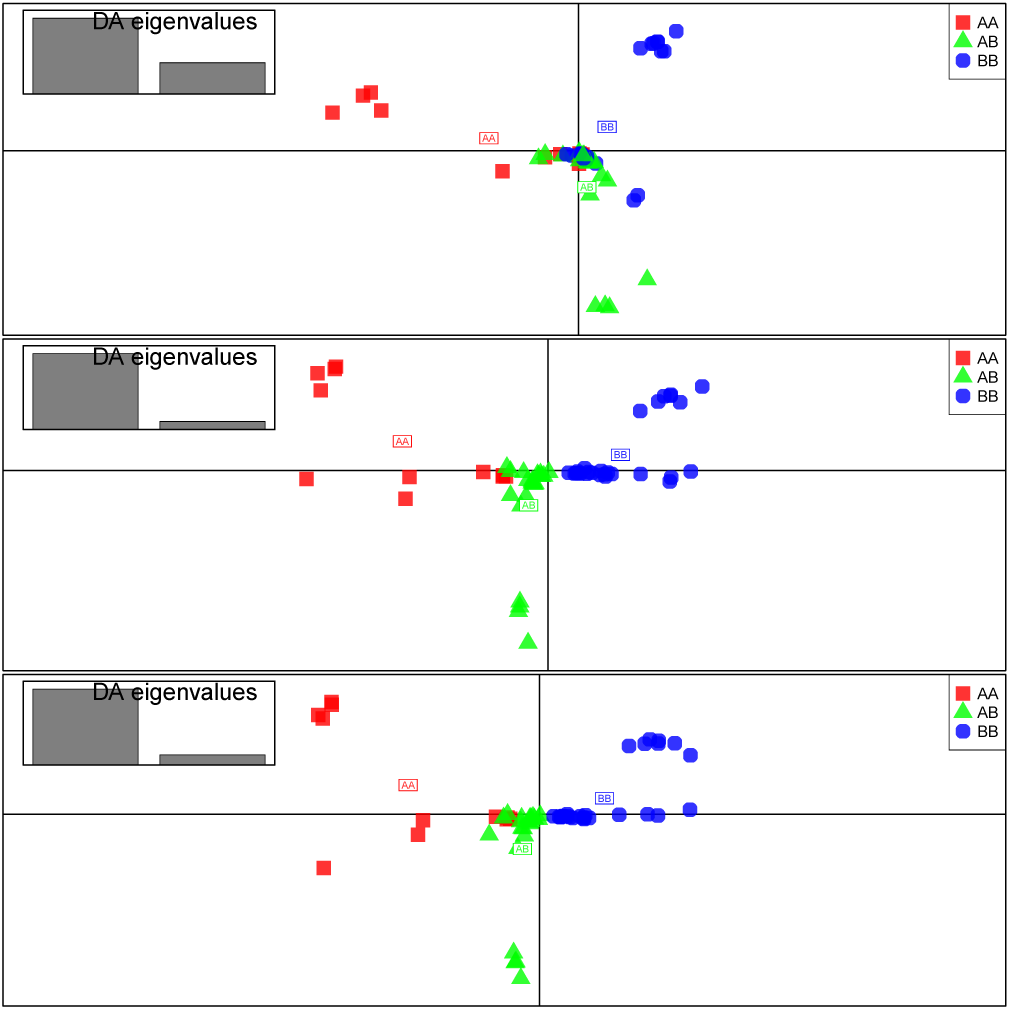
Discriminant analysis of principal components (DAPC) of genomic variation among ecotypes of Icelandic cod. Variation of genomic regions that can be mapped to linkage groups LG02 to LG23 (excluding LG01) (top panel), mapped to all linkage groups (middle panel), and the rest of the genome that cannot be mapped to linkage groups (bottom panel). DA, retaining 15 principal components by cross-validation, was based on *Pan* I genotypes (*AA, AB, BB*) of LG01 that were used as a proxy for behavioural ecotypes.

Our interpretation of the totality of the genomic evidence (based on up to 8.6 million variable sites after filtering) was that the shallow-water coastal (some *AA*) and deep-sea frontal (some *BB*) ecotypes are reproductively isolated coastal and frontal species. They are cryptic species within the Atlantic cod complex that have adapted to environmental factors in shallow and deep waters and diverging at LG01. They hybridized and formed a new homoploid hybrid species. There is normal Mendelian segregation of variants within the true-breeding hybrid species (such as the *Pan*I *AA*, *AB*, and *BB* on LG01). This addresses the contradictory results regarding intermediate forms^23^ and has implications for the interpretation of the relative importance of these groups (e.g., the contribution of the different Greenland ecotypes to the Icelandic population^31,35^ and vice versa).

The hybrid individuals defined by posterior membership probabilities shown in the genomic results (Supplemental Figure 6) were on average intermediate regarding phenotype and habitat use between the two pure types (Supplemental Figure 7). However, hybrids were also more variable and transgressed the phenotype of the parental forms.^1,2^ Considerable variation was observed regarding behaviour, as determined using storage-tag data, among individuals classified by the *Pan* I genotype. There were *AA* individuals that showed a frontal behaviour and *BB* individuals that exhibited a coastal behaviour (see appendix in^23^ ). This behavioural variation may reflect species differences within the *Pan* I genotypes. This heterogeneity of behavioural types provides independent support for our thesis.

It was difficult to reconcile the long-distance linkage disequilibrium^8^ and the behaviour of genomic islands observed here (for example with certain genomic islands specific to a particular location) with a model of speciation with gene flow.^36^ A modified model of divergence after speciation^36^ explains inter-chromosomal correlations and is applicable to this case of homoploid hybrid speciation. Under this model we assume rapid speciation by hybridization of already divergent lineages (Figure 4), possibly involving chromosomal rearrangements^11^ and further divergence after speciation. The large genomic islands are islands of ecological adaptation^20^ and/or sites of low recombination,^36^ and not necessarily islands of speciation. Speciation genes may reside elsewhere in the genome.^36^

## Conclusions

Our study demonstrates the power of whole-genome sequencing and population genomics in providing deep insights into fundamental processes of speciation. The results raise many interesting hypotheses about the extent of hybrid speciation and its role in cryptic biodiversity in the sea.^4^ Our results also raise hypothesis about marine fish in general and about codfish biology in particular, which remain to be tested. For example, we need to examine the criteria of hybrid speciation^10^ and hybrid sterility, as well as structural chromosomal variation,^11^ age of hybridization, biogeography and natural selection and adaptation to diverse environments such as deep and shallow^27^ or low salinity^37^ waters. Our results call for the re-evaluation of previous work on cod-fish with implications for resource management. The hybrid nature of both the walleye pollock and Atlantic cod raises the question concerning the extent to which very profitable fisheries^12,13^ depend on hybrid vigour. For example, did the decimation of the hybrid and frontal fish influence the collapse and non-recovery of the western fisheries^38^? Ocean changes and their impact of ecosystems and fisheries have unpredictable consequences.^14^ The ongoing climate warming and northern hemisphere ice melting predict species interchange between the north Pacific and the Atlantic.^15^ The potential exists for further hybridization of Pacific, Arctic and Atlantic taxa, with unknown consequences for biodiversity.

## Methods

**Population sampling** We randomly sampled over 200 individual cod-fish from our large sample collection of greater than 20,000 Atlantic, Arctic and Pacific cod-fish individuals.^39,40^ We stratified the sampling to cover the widest geographic range possible with our database. We sampled Pacific cod *Gadus macrocephalus* (Tilesius, 1810) (mnemonic: Gma) and walleye pollock *Gadus chalcogrammus* (Pallas, 1814) (Gch) from the Pacific Ocean. We sampled Polar cod *Arctogadus glacialis* (Dryagin, 1932) (Agl), Arctic cod *Boreogadus saida* (Lepechin, 1774) (Bsa), and Greenland cod (uvak) *Gadus ogac* (Richardson, 1836) (Gog) from the Arctic at western Greenland. We sampled Atlantic cod *Gadus morhua* (Linnaeus, 1758) (Gmo) from throughout its distribution in the North Atlantic Ocean. The localities range west to east from off Chatham on Cape cod (mnemonic Cco), the Western Bank (Web) and Sable Bank (Sab) off Nova Scotia, Trinity Bay (Tri) and the Southern Grand Banks (Sgb) of Newfoundland. Collectively we called these localities the West. We sampled from the west and east coast of Greenland (Gre), around Iceland (Ice), the Barents Sea (Bar), the White Sea (Whi), and coastal Norway (Nor). Collectively we called these localities the North. Furthermore we sampled from the Faroe Islands (Far), the North Sea (Nse), the Celtic Sea (Cel), and the Western and Eastern Baltic (Bal-W and Bal-E respectively). Collectively we called these localities the South. The sample from Iceland included a number of individuals of the three genotypes of the *Pan* I locus (*AA*, *AB*, and *BB*)^25^ located on linkage group LG01^26^ which we used as a proxy to identify behaviourally consistent ecotypes of coastal, intermediate, and frontal cod as defined by results from storage-tags data.^23,24,28^

The molecular and morphological relationship and biogeography of these taxa have been discussed^32,41,42^ and the the most comprehensive account is based on mitochondrial genomics.^5^ Coulson et al^5^ consider Arctic cod (Bsa) to be an outgroup for all these taxa. Atlantic cod (Gmo) and walleye pollock (Gch) are the most closely related taxa and Pacific cod (Gma) slightly more distant. They argue^5^ that Pacific cod and walleye pollock represent two separate but nearly simultaneous invasions of the Pacific Ocean. They date the Atlantic cod vs Pacific cod split at 4 Mya and the Atlantic cod vs walleye pollock split at 3.8 Mya using conventional rates of mtDNA evolution. They^5^ suggested a nomenclature revision from *Theragra chalcogramma* to *Gadus chalcogrammus* (Pallas, 1814) for walleye pollock that has been accepted by the American Fisheries Society.^43^ We follow the new nomenclature here. According to their view^5^ Greenland cod (Gog) is a recent reinvasion of Pacific cod (Gma) into the Arctic and they consider it to be a subspecies of Pacific cod. Their analysis is based on a hypothesis of speciation by lineage splitting. We designed our sampling partly to investigate these relationships.

**Molecular analysis** We prepared samples for high-coverage (20 – 30×) and for low-coverage (2×) sequencing on the Illumina HiSeq 2500 platform. We randomly selected one individual each of Arctic cod (Bsa), Pacific cod (Gma), walleye pollock (Gch), and Greenland cod (Gog) for a 20× sequencing coverage. We randomly selected six Atlantic cod (Gmo), two from the West, two from Iceland, and two from the North Sea, for 30× sequencing coverage. For the low-coverage analysis we randomly selected about a dozen individuals of Arctic cod (Bsa), Pacific cod (Gma), walleye pollock (Gch), and Greenland cod (Gog), one individual of Polar cod (Agl), and 152 Atlantic cod (Gmo) individuals from various localities.

Our tissue collection is primarily gill tissue but also has fin clips and muscle tissue and DNA isolated from blood.^44^ Tissues are stored in 96% ethanol. We isolated genomic DNA from 10 individuals selected for high-coverage sequencing using the NucleoSpin^®^ Tissue kit (Machery-Nagel, reference 740952.50) following the manufacturer’s protocol. We isolated genomic DNA from 200 individuals selected for low-coverage sequencing using the E.Z.N.A.^®^ Tissue DNA Kit (Omega biotek) following the manufacturer’s protocol.

We quantified and estimated 260/280 and 260/230 quality of the genomic DNA using a NanoDrop 2000 UV-Vis Spectrophotometer (Thermo Scientific). We also quantified genomic DNA using fluorescent detection with the Qubit^®^ dsDNA HS Assay Kit (Life Technologies).

Libraries for the high-coverage sequencing were made at the Bauer Core Facility at Harvard University. The facility used the Covaris S220^®^ (Covaris) to shear the genomic DNA to a target size of 550 bp. The Apollo 324^TM^ (Wafergen Biosystems) system was used to generate libraries for DNA sequencing. Each library had a single index. The size distribution of the libraries was determined with the Agilent Bioanalyzer (Agilent Technologies). Library sample concentration was determined with qPCR according to an Illumina protocol.

We prepared libraries for low coverage sequencing using the Nextera^®^ DNA Library Preparation Kit (Illumina, FC-121-1031). We used the Nextera^®^ Index Kit (FC-121-1031) with dual indices. Index N517 was used instead of N501. We followed the manufacturer’s Nextera protocol. We cleaned the tagmentated DNA with a Zymo Purification Kit (ZR-96 DNA Clean & Concentrator^TM^ -5, Zymo Research). We cleaned PCR products and size selected with Ampure XP beads (Beckman Coulter Genomics, reference A63881). We used the modification recommended for PCR clean-up for 2×250 runs on the MiSeq using 25*μ*l of Ampure XP beads (instead of 30*μ*l) for each well of the NAP2 plate. We quantified the individual libraries with fluorescent detection using a Quant-iT^TM^ PicoGreen^®^ dsDNA Assay Kit Quantit kit on a Spectramax i3x Multi-Mode Detection Platform (Molecular Devices). We determined the size distribution of 12 randomly chosen libraries. We then normalized the multiplexed DNA libraries to 2nM concentrations and pooled the libraries. The size distribution of the pooled libraries was determined by the Bauer Core Facility using the Agilent Bioanalyzer. Pooled library sample concentration was determined with qPCR.

Pooled libraries were sequenced on the HiSeq 2500 in rapid run mode (paired-end, 2×250 cycles) at the Bauer Core Facility at Harvard University.

**Statistical analysis** The Bauer Core Facility through the department of Informatics and Scientific Applications returned the base-called data as de-multiplexed `fastq` files. Individuals were sequenced on two lanes of the HiSeq 2500. We merged both of the forward and the reverse reads for each individual. We fetched the `Gadus_morhua.gadMor1.dna.toplevel.fa` genomic reference sequence from www.ensemble.org and used it as a reference sequence. We aligned the merged `fastq` reads to the reference using `bwa mem`.^45^ We used `samtools` to generate sorted and indexed bam files from the `sam` files.

Next generation sequencing data of this kind has high error rates from multiple sources, including base-calling and alignment errors.^9^ The low coverage data suffer from these errors in particular. Such errors will affect downstream analysis that depend on calling SNPs and genotypes because errors will be compounded. We adopted the strategy of using methods based on genotype likelihoods^9^ implemented in the `ANGSD`^46^ and related software.^47–51^ These methods yield SNP and genotype information and an associated uncertainty facilitating unbiased or low-biased statistical interpretation of low-coverage data. Using these tools allowed us to sequence a larger number of individuals.

We used `NGSadmix`^51^ to estimate individual admixture proportions based on a model of *k =* 2…*n* ancestral populations. We used `ngsDist`^50^ to estimate pairwise genetic distances and used `fastME` (version 2.07^52^) to make minimum evolution phylogeny from these distances. We used `ngsCovar`^50^ to estimate a covariance matrix for principal component analysis (PCA) and discriminant analysis of principal components (DAPC).^53^

The analysis is based on not calling genotypes or alleles but instead using genotypic likelihoods^9^ as implemented in the ANGSD software^46^ for fast analysis of large samples. We used `realSFS` of `ANGSD`^46^ to estimate site frequency spectra and *F_ST_*. We used `ANGSD` and friends for quality filtering. We typically used a minimum mapping quality of 30 and minimum base quality of 20 `(–minMapQ 30 –minQ2 0)` and discarded bad reads `(–remove_bads 1)`, filtered out sites with a minor allele frequency less than 0.05 `(–minMaf 0. 05)`, filtered by number of individuals `(e.g. –minInd 10)`, and filtered out sites that are very likely to be polymorphic with a *P* value less than 10^−6^ `(–SNP_pval 1e–6)`. We did a pairwise sliding-window *F_ST_* analysis with a window size of 10,000 and a step size of 2,000.^46^ As an example of the effects of filtering the admixture analysis of Atlantic cod had 22.6 million sites that were reduced to 8.6 million sites after filtering.

We used R^54^ and various `Rscripts` from the `ANGSD` and friends packages and our own functions and scripts to manipulate and plot the results. We did Principal Component Analysis (PCA) using the `eigen` function in `R` and plotted results with `ggplot2`.^55^ We used `adegenet`^53^ for Discriminant Analysis of Principal Components (DAPC). We performed cross-validation (CV) using the `xvalDapc` function^53^ to determine the number of principal components (PCs) to retain in the DAPC. We used the posterior membership probabilities returned by DAPC of the whole-genome sequence data to define “species” as priors for a DAPC of phenotypic and habitat use data.

Large ’genomic islands of divergence’ in the Atlantic cod genome^6,7^ are already known. The divergence of these islands may influence analysis of genome wide effects. To address these problems we separately analysed the parts of the genome that can be mapped to specific linkage groups as well as the parts that cannot be mapped to linkage groups. In order to map Atlantic cod genomic scaffolds^56^ to the Atlantic cod genetic map^26,57^ we did a local `blastn`^58^ of the 120 base pairs (bp) surrounding the SNPs used for mapping^26^ onto the reference sequence and chose matches that were both greater than 100 bp and greater than 95% identity. The genetic map^26,57^ is sparse and about 235 megabase (Mb) (roughly 1/3 of the genome) can be thus assigned to specific linkage groups. We refer to the parts of the genome that map to linkage groups LG02 to LG23 as that part of the genome which definately does not map to linkage group LG01. We then add LG01 data and refer to it as the part of the genome that is mapped to linkage groups. We also analyse the parts of the genome (about 2/3) that cannot be thus mapped to specific linkage groups.

The computations in this paper were run on the Odyssey cluster supported by the FAS Division of Science, Research Computing Group at Harvard University.

## Acknowledgements

We thank Hopi Hoekstra for laboratory space, Kyle Turner for technical help, John Wakeley for help with computing resources, and Rasmus Nielsen for help on design. We thank Karsten Hartel, Andrew Williston and Ubaldo Benitez Hernandez for help with specimen transport. We thank Grant Pogson, Michael Canino, Jarle Mork and Krisján Kristins-son for help with sampling. We also especially thank Dick Lewontin who made it all possible.

The work was supported by a grant from Svala Árnadóttir private fund, by a grant from the University of Iceland Research Fund, and by institutional funds from R.C. Lewontin.

## Author contributions statement

E.Á. and K.H. conceived the experiments, E.Á. and K.H. conducted the experiments, E.Á. and K.H. analysed the results. Both authors wrote and reviewed the manuscript.

## Additional information

**Accession codes:** SRP065670. We request that users of the data honor Fort Lauderdale data sharing principles of granting us first publication of results from our genomic data.

**Competing financial interests:** The authors declare no competing financial interests.

## Supplemental Information

**Supplemental Figure 1.**
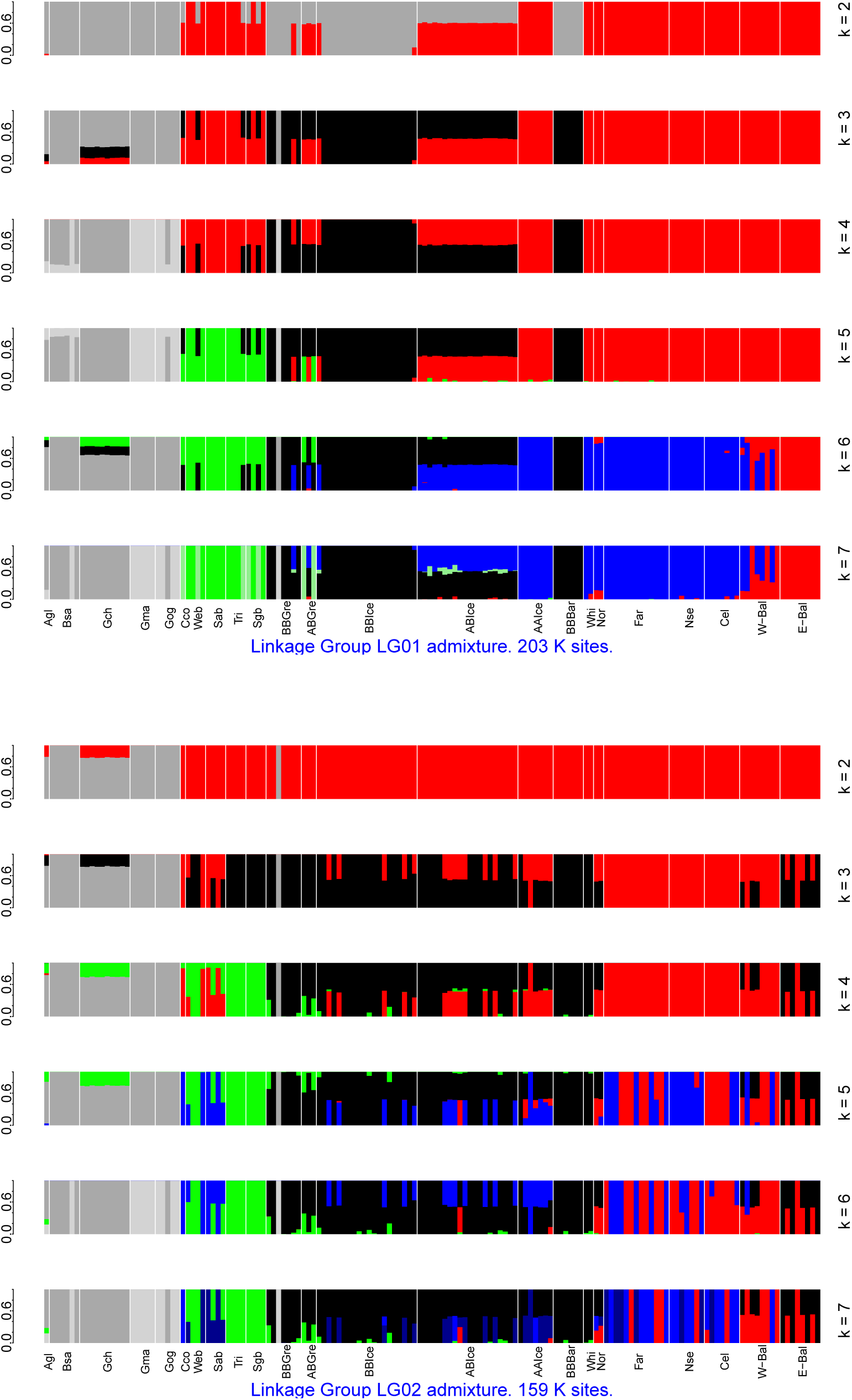
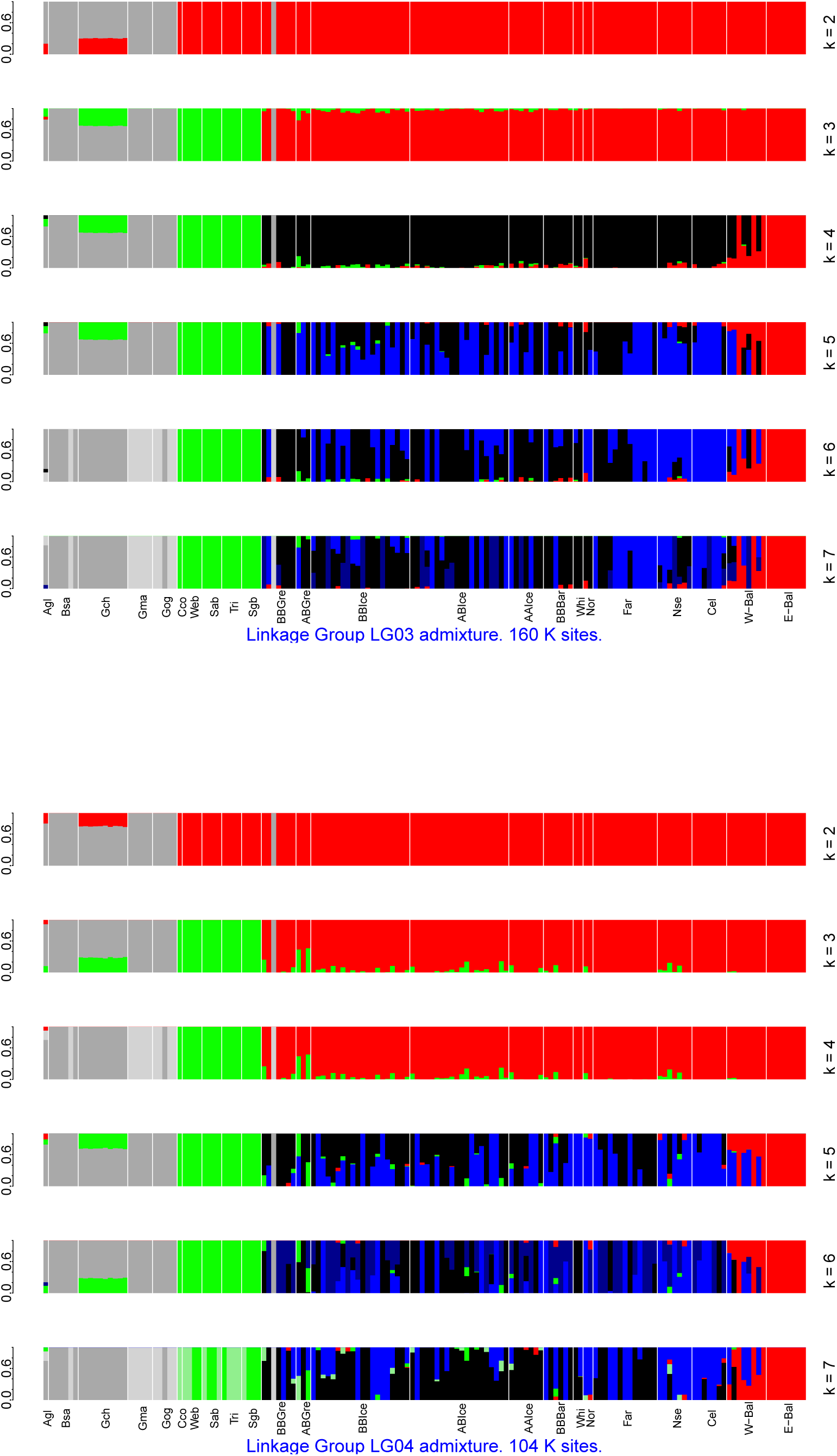
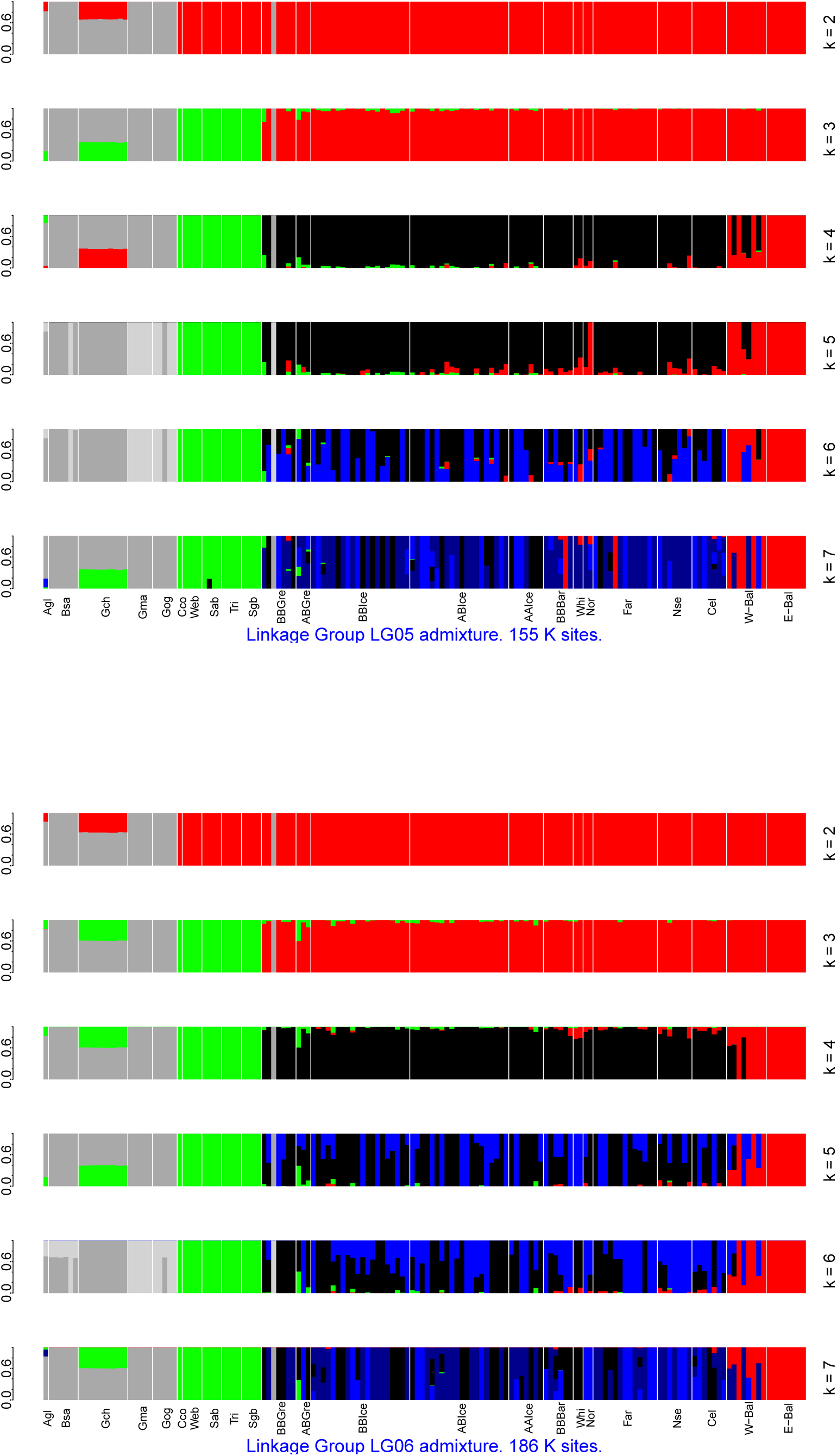
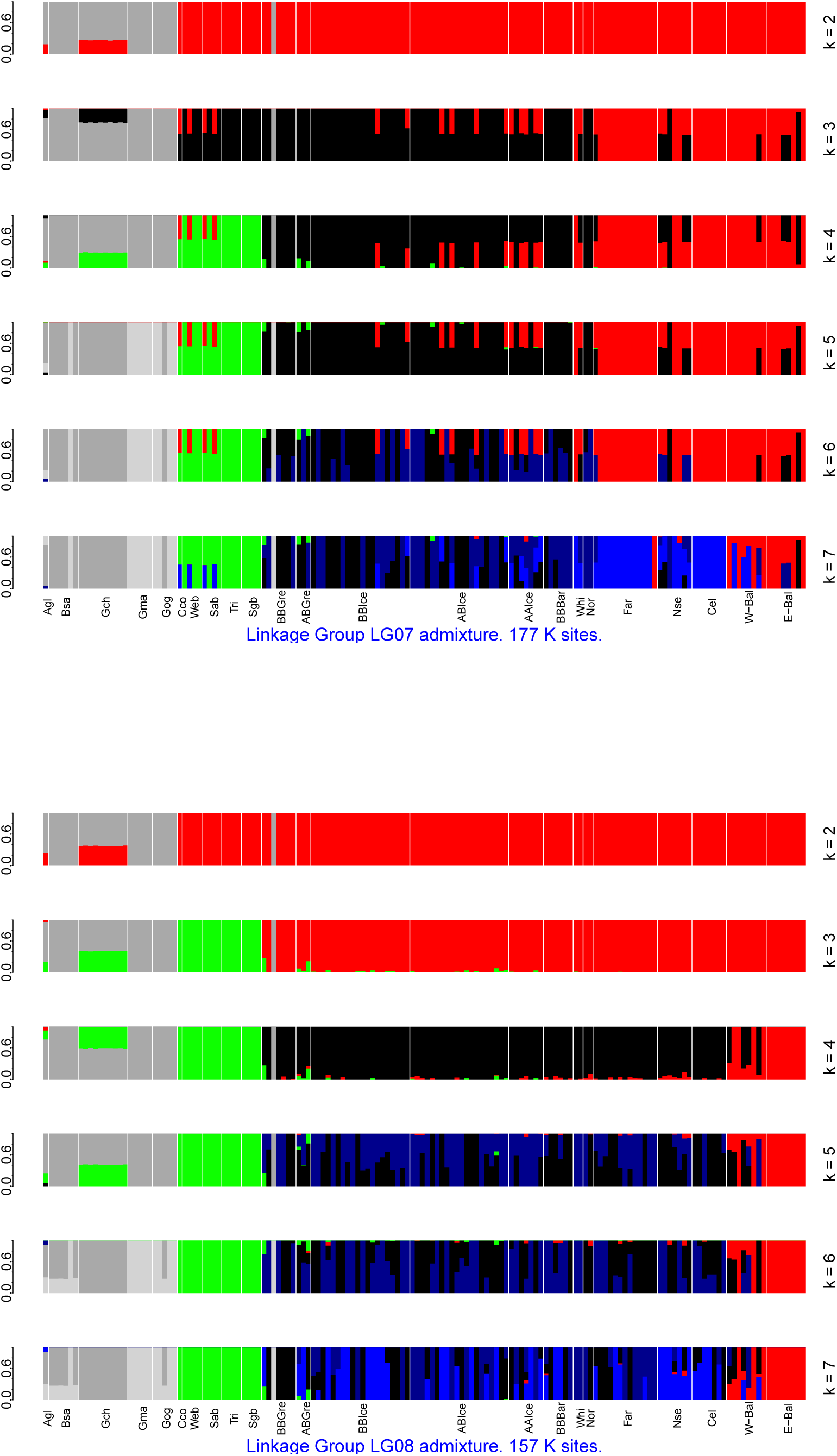
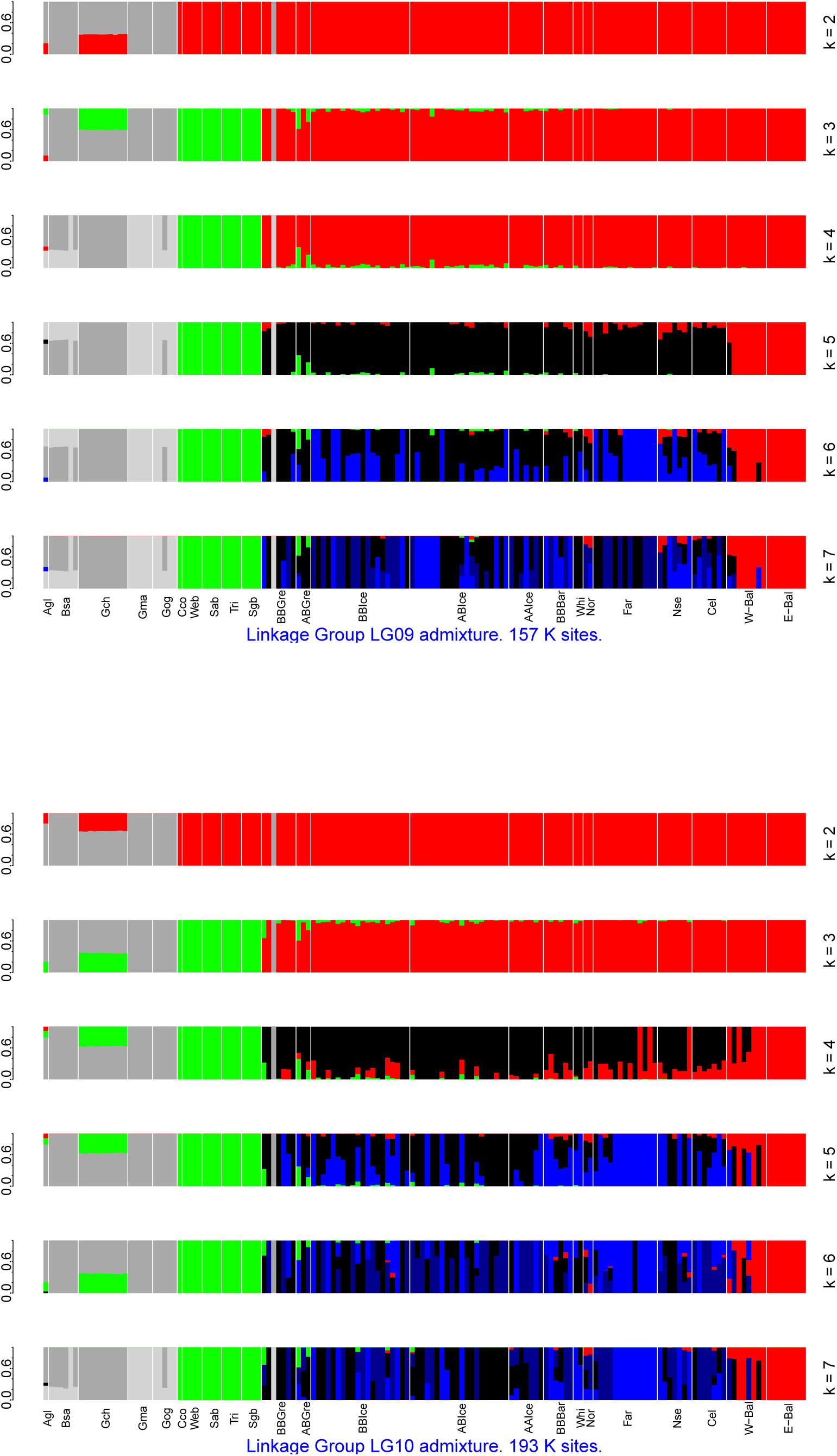
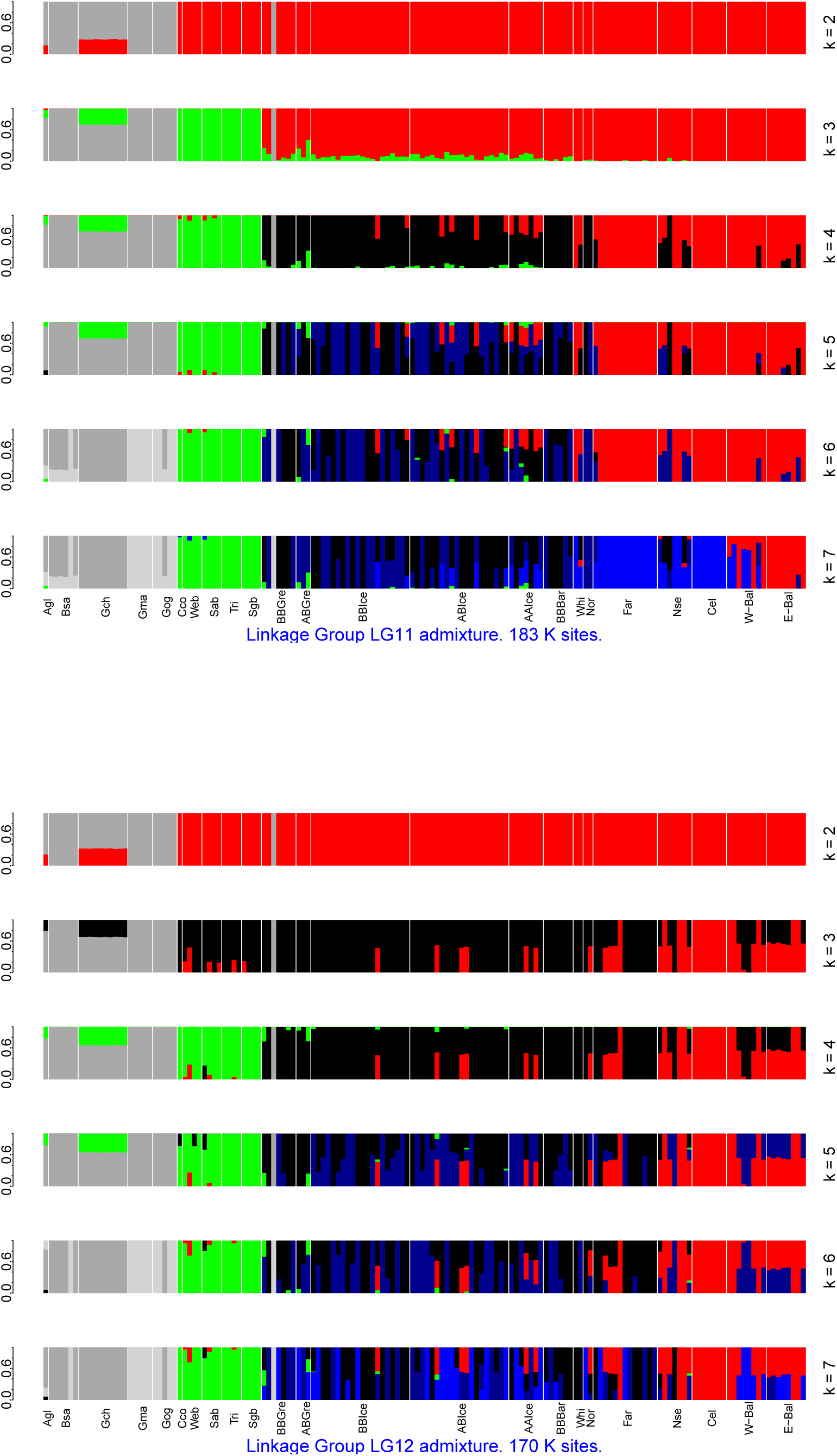
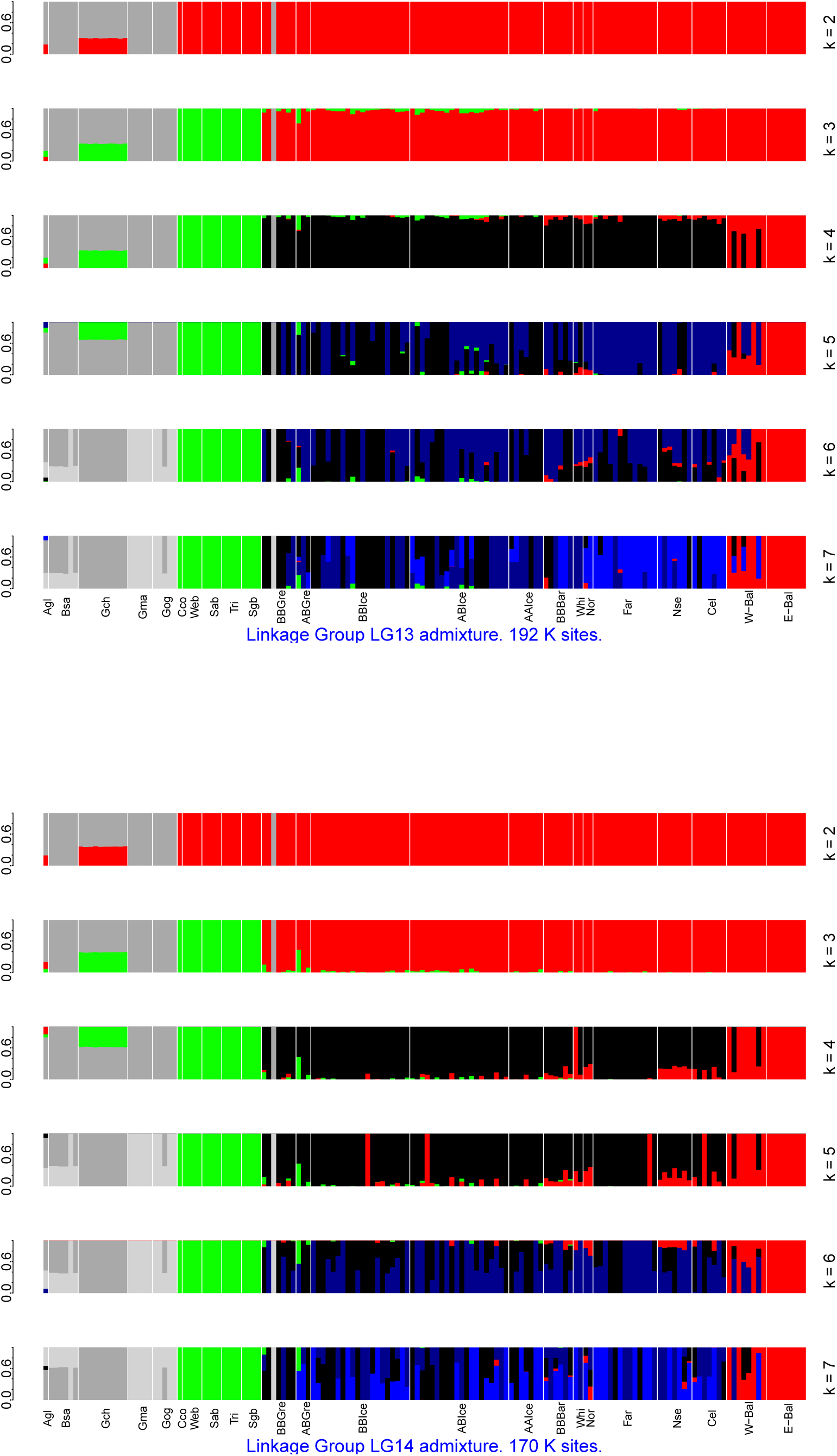
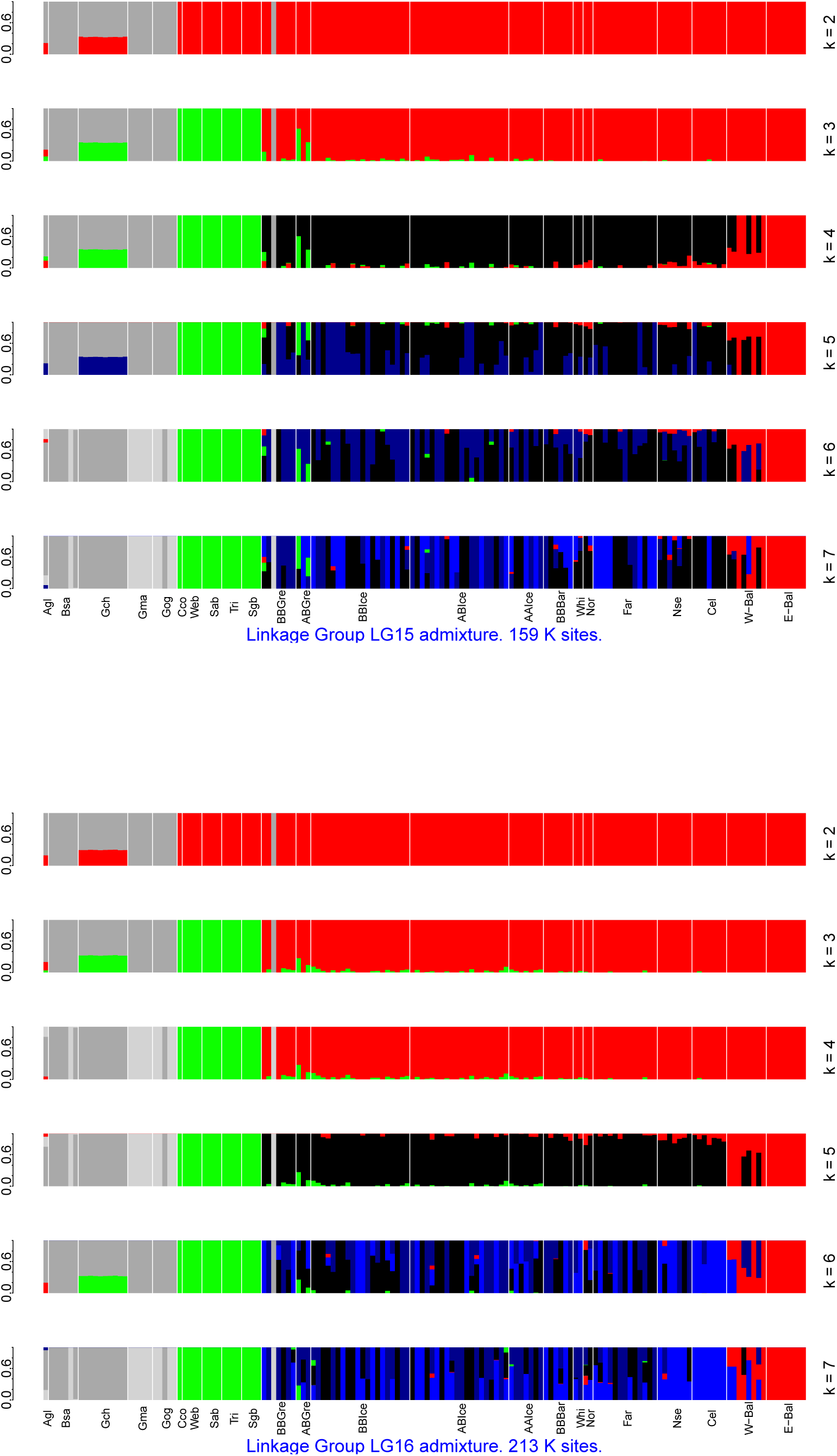
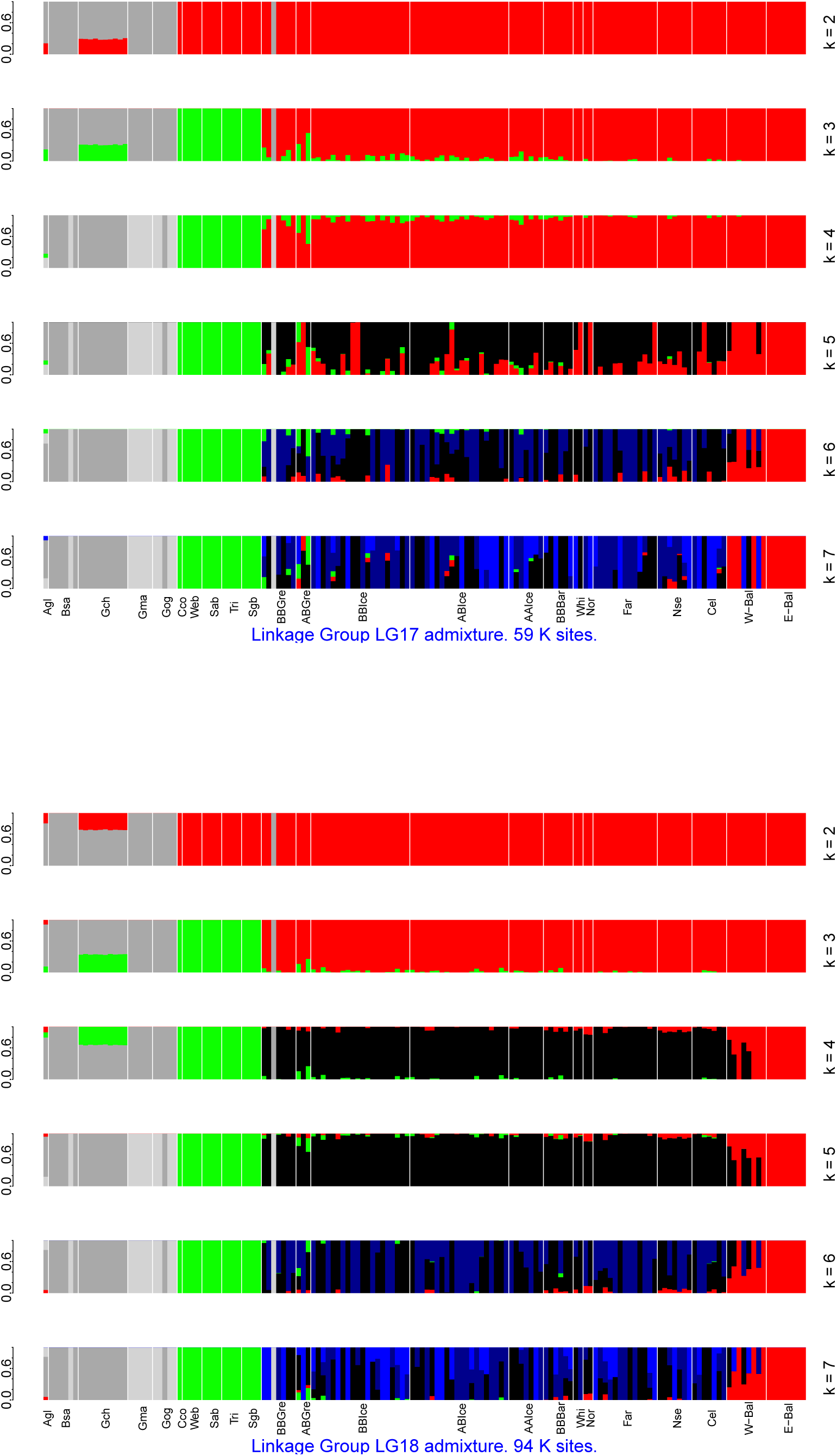
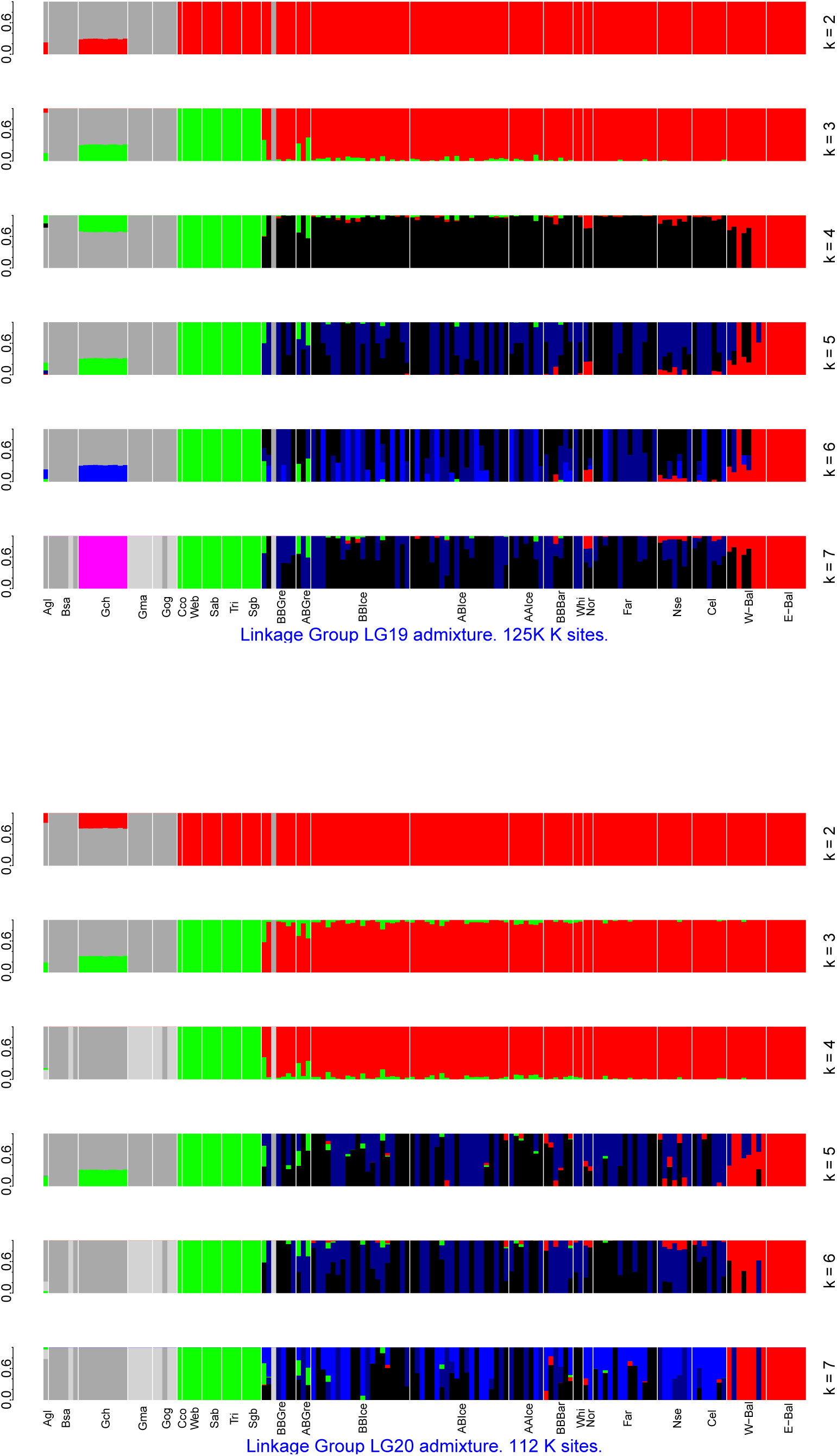
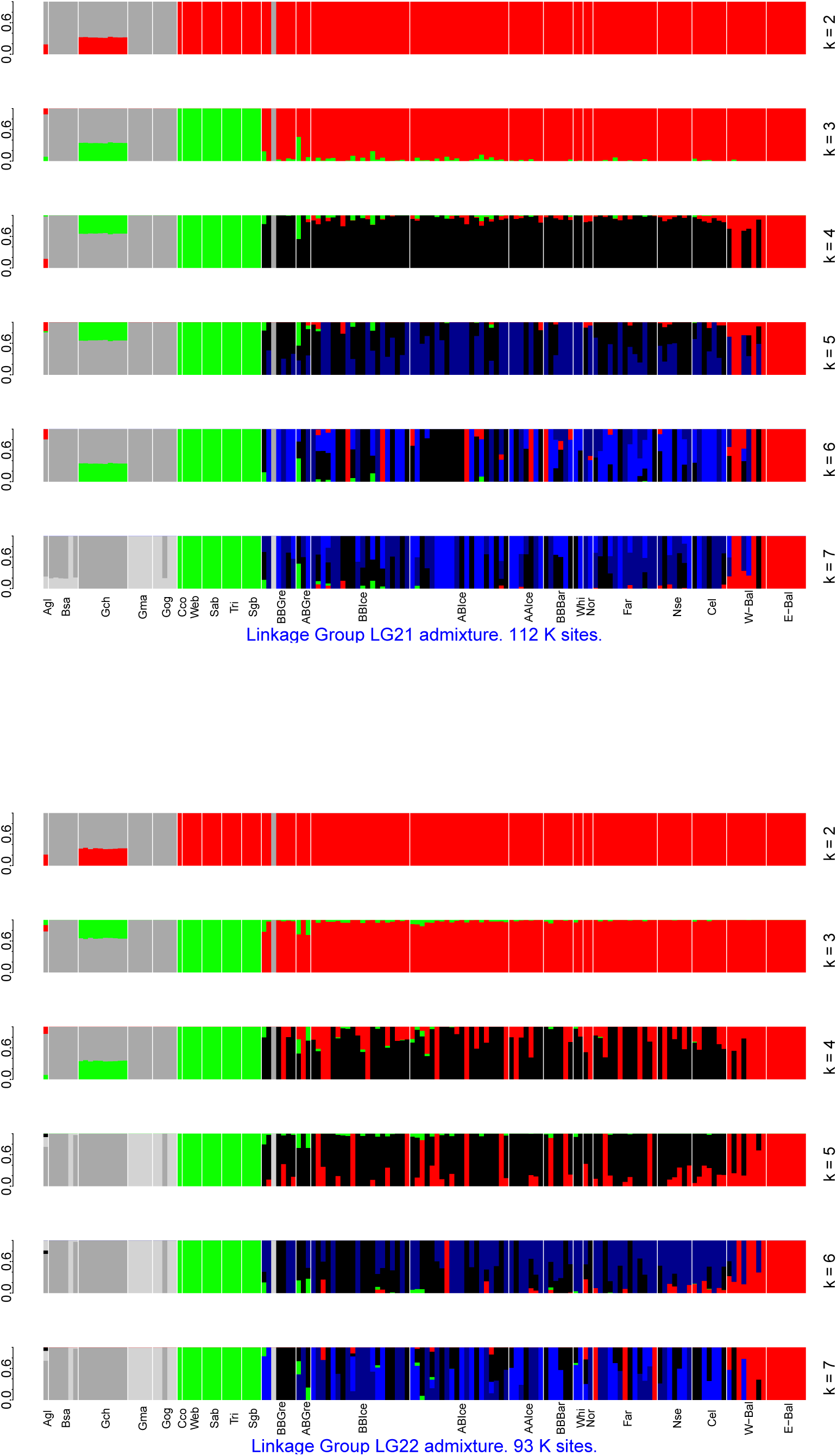
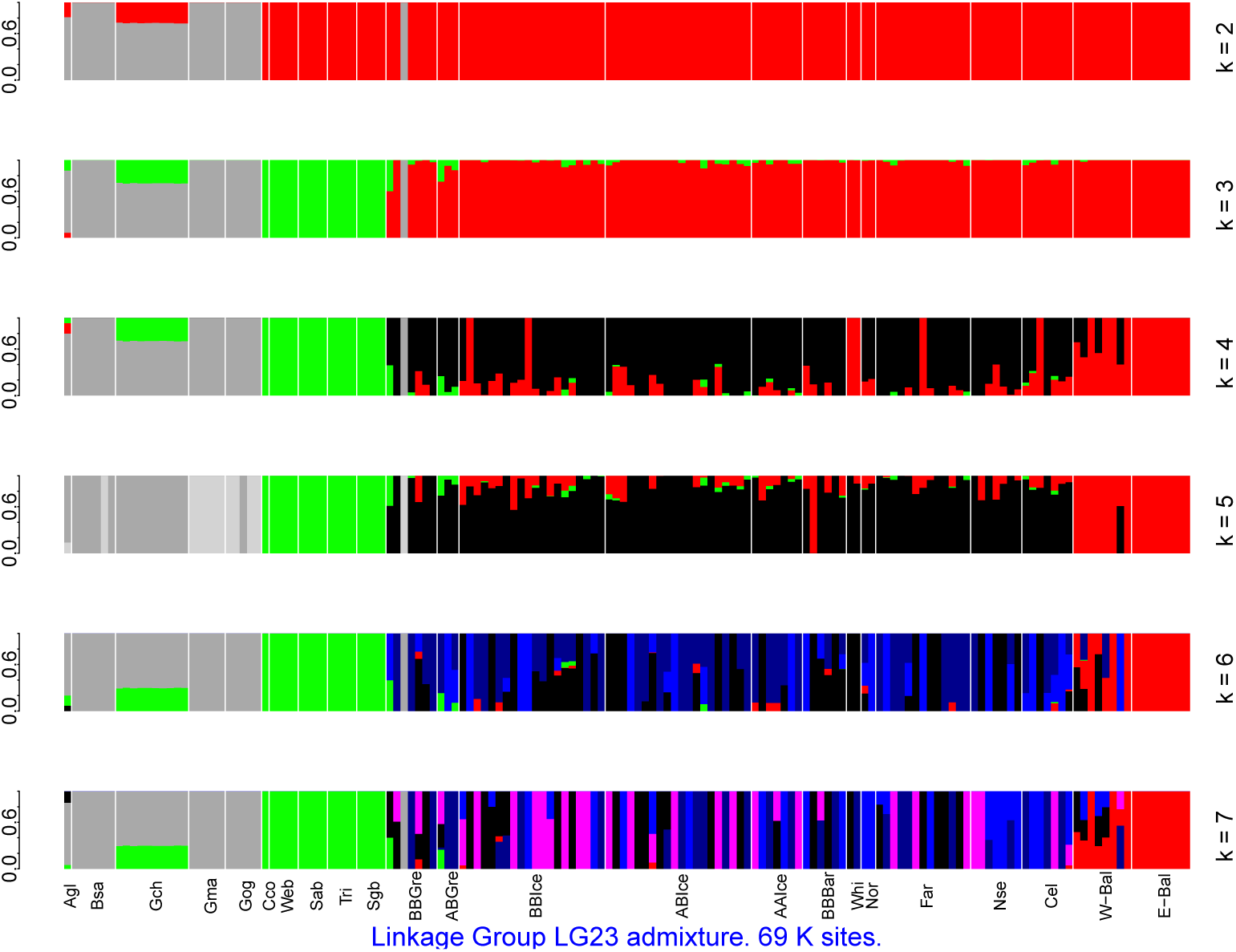
Individual admixture analysis of cod-fish by linkage group LG01 to LG23. Model-based analysis with the number of ancestral population *k* ranging from 2 to 7. Ordinate gives admixture proportion of each individual. Results ordered based on species *Arctogadus glacialis* Agl, *Boreogadus saida* Bsa, *Gadus chalcogrammus* Gch, *Gadus macrocephalus* Gma, *Gadus ogac* Gog, and for *Gadus morhua* localities from west to east and by ecotype: Cape cod Cco, Western Bank Web, Sable Bank Sab, Trinity Bay Tri, Southern Grand Banks Sgb, Greenland frontal BBGre, Greenland intermediate ABGre, Iceland frontal BBIce, Iceland intermediate ABIce, Iceland coastal AAIce, Barents Sea frontal BBBar, White Sea Whi, Norway Nor, Faroe Islands Far, North Sea Nse, Celtic Sea Cel, western Baltic W-Bal, and eastern Baltic E-Bal. The number of variable sites in thousands (K) is given for each linkage group.

**Supplemental Figure 2.**
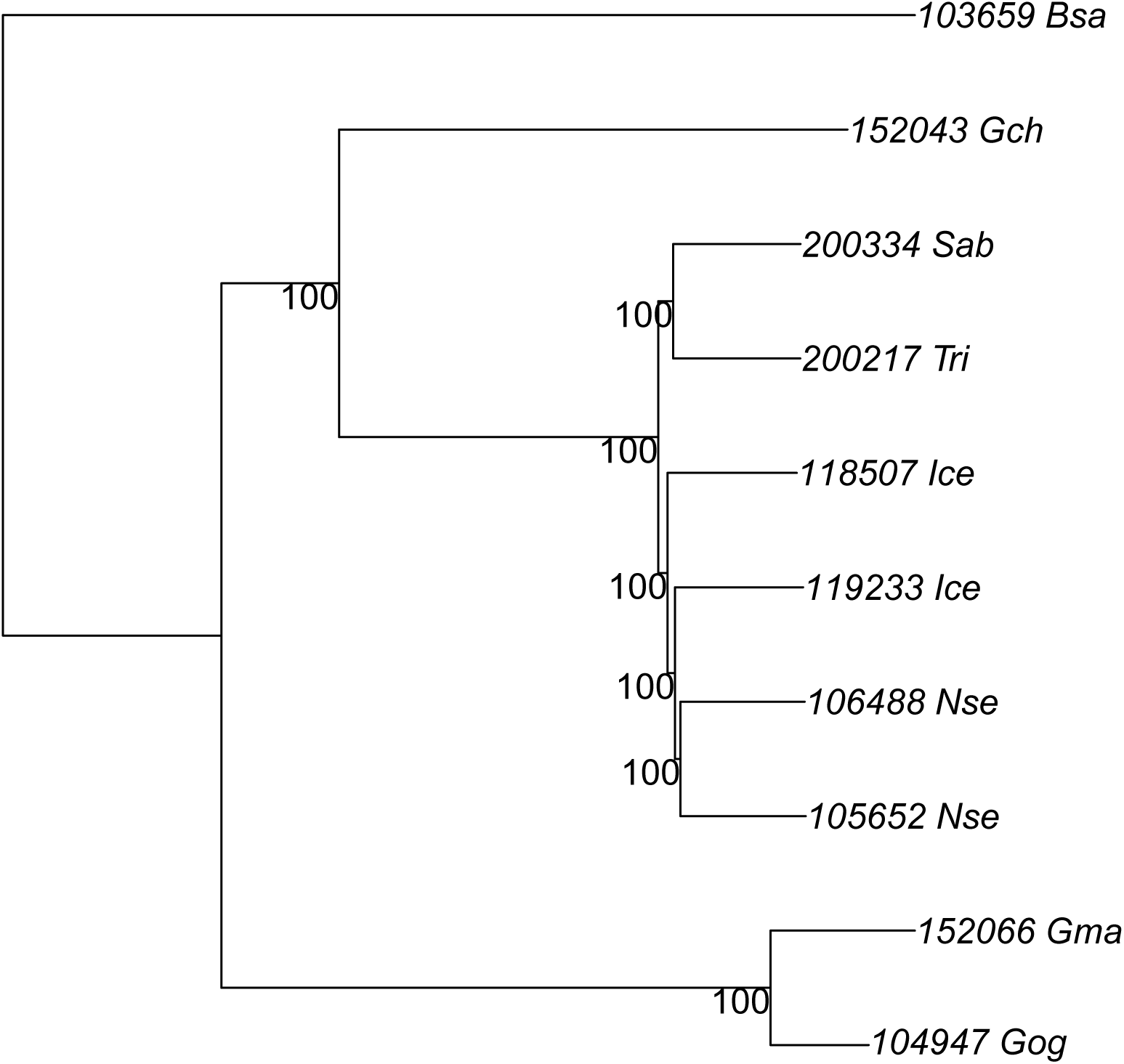
Minimum evolution tree of genetic distances among cod-fish taxa. *Boreogadus saida* Bsa, *Gadus chalcogrammus* Gch, *Gadus macrocephalus* Gma, *Gadus ogac* Gog and six *Gadus morhua* Gmo, from Sable Bank Sab, Trinity Bay Tri, Iceland Ice, and the North Sea Nse. Based on whole genome sequencing with 20 – 30× coverage of entire genome. Individuals are labelled with a six digit identifier plus a three letter code for species or localities for Atlantic cod. The numbers on the nodes are bootstrap supports.

**Supplemental Figure 3.**
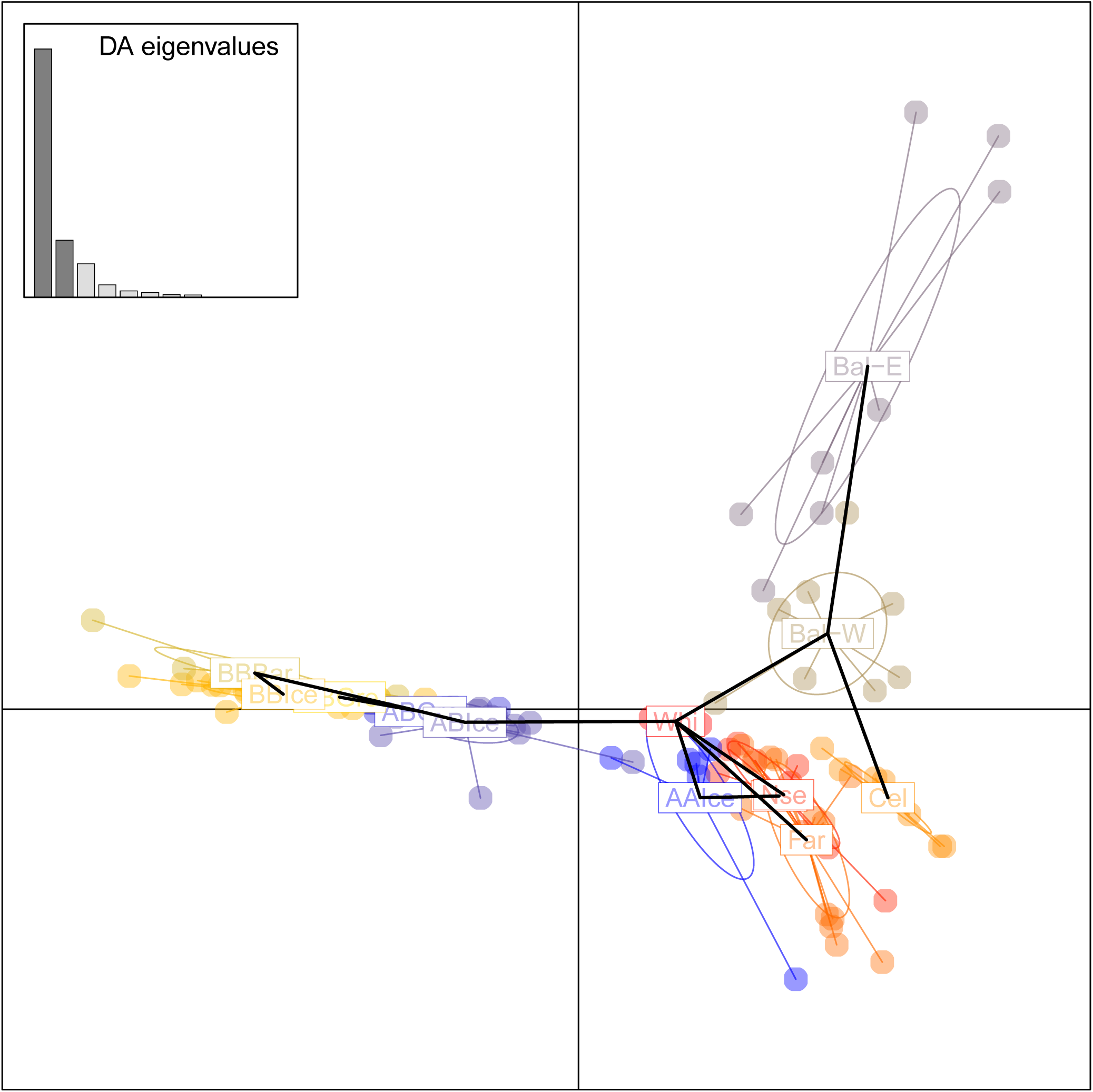
Discriminant analysis of principal components (DAPC) of variation among groups and localities of the eastern Atlantic. Black lines represent a minimum spanning tree. Variation of the part of the genome mapping to linkage groups LG01 to LG23. DAPC based on deep sea frontal fish from the Barents Sea (BBBar), Iceland (BBIce), and Greenland (BBGre), intermediate fish from Iceland (ABIce) and Greenland (ABGre) and finally shallow water coastal fish from the White Sea (Whi), Iceland (AAIce), Faroe Islands (Far), Norway (Nor), North Sea (Nse), Celtic Sea (Cel), and the western and eastern Baltic (W-Bal and E-Bal).

**Supplemental Figure 4.**
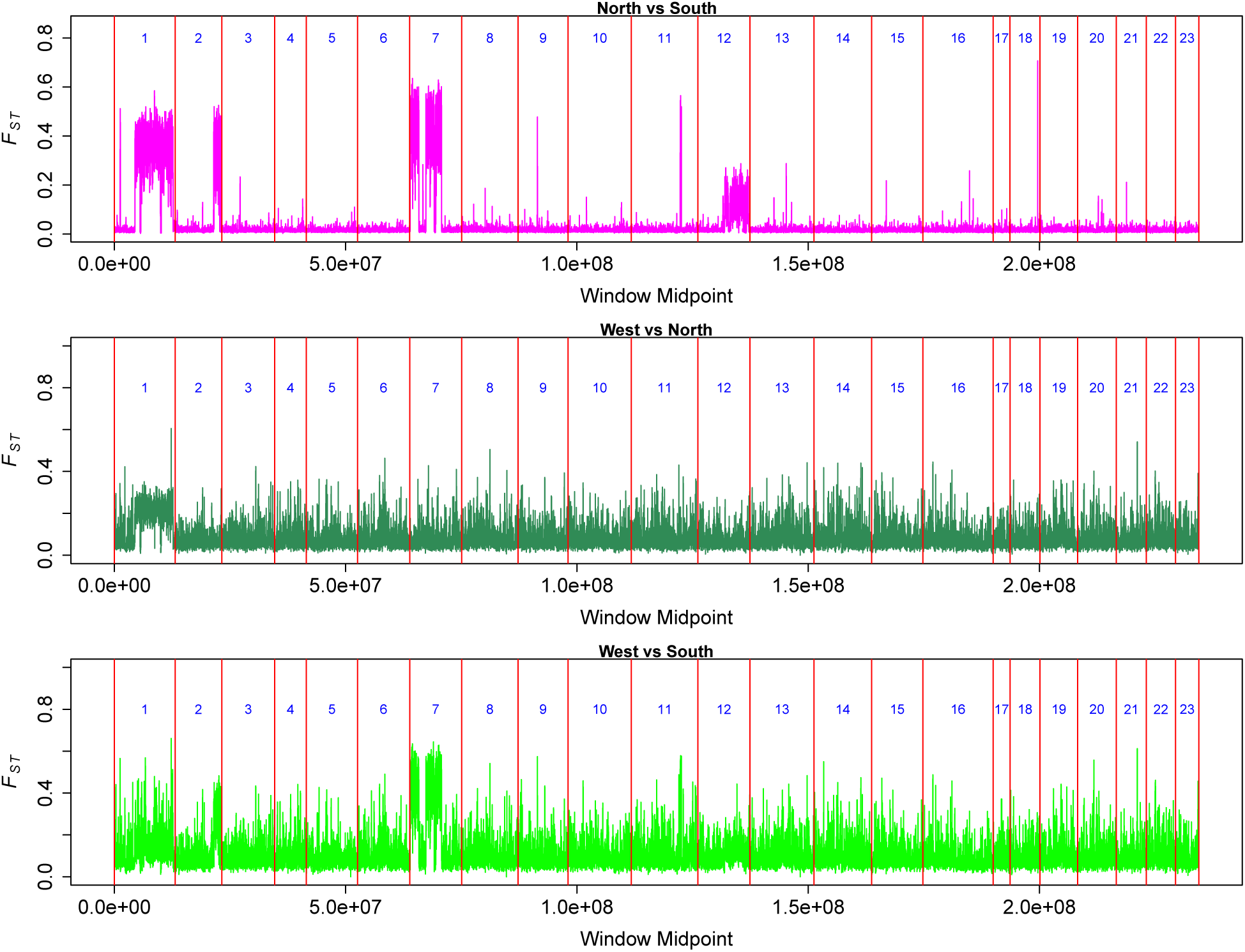
Sliding-window *F_ST_* on window midpoint. Genomic data that map to linkage groups LG01 to LG23 among localities of Atlantic cod. Pairwise comparisons of West (Cape Cod, Western Bank, Sable Bank, Trinity Bay, and Southern Grand Banks pooled), North (Greenland, Iceland, Barents Sea, White Sea, and Norway pooled), and South (Faroe Islands, North Sea, Celtic Sea, and Baltic Sea pooled). Window size was 10,000 and step size was 2,000.

**Supplemental Figure 5.**
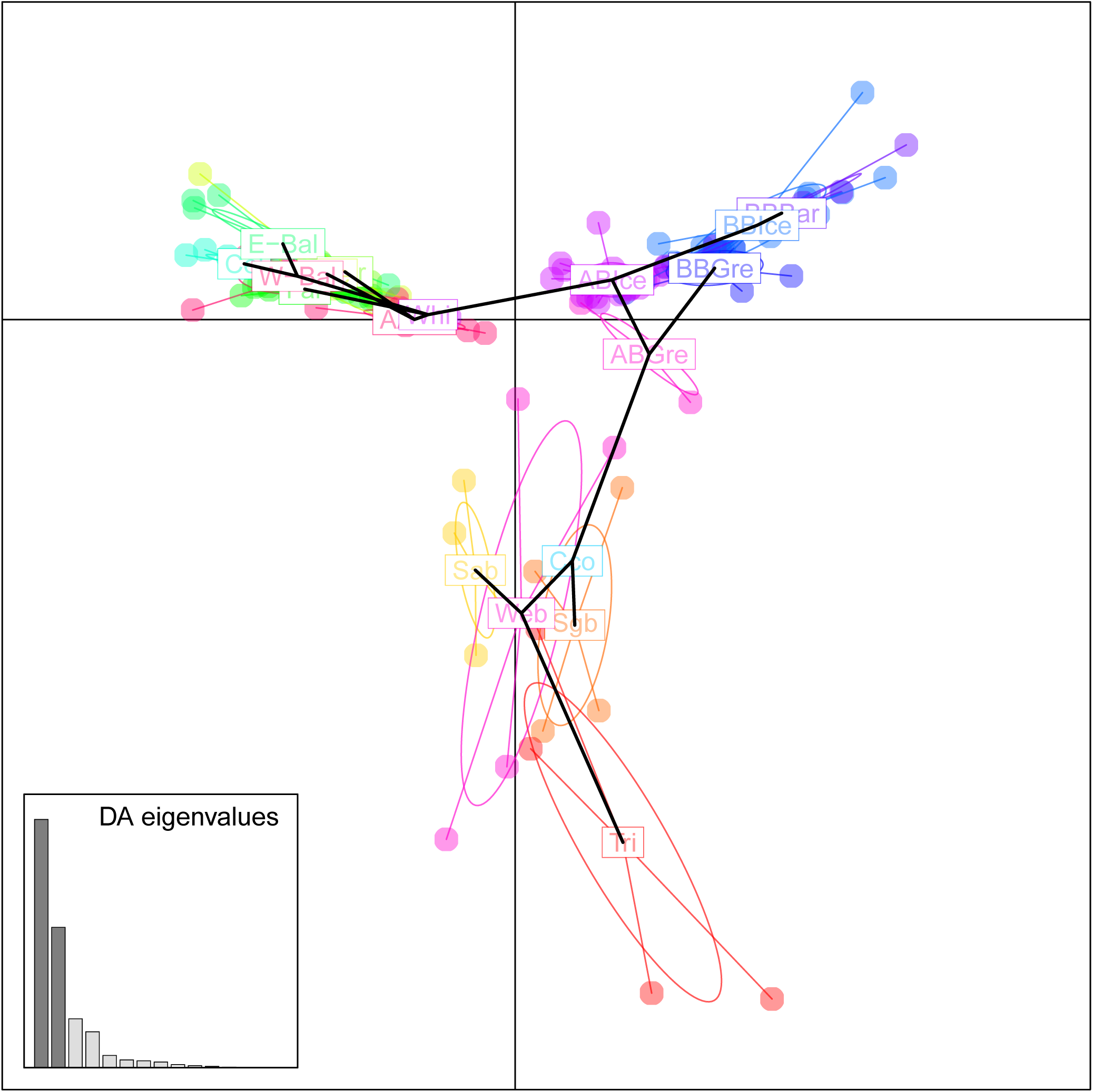
Discriminant analysis of principal components (DAPC) of variation among groups and localities of Atlantic cod. Black lines represent a minimum spanning tree. Variation of the part of the genome mapping to linkage groups LG01 to LG23. DAPC based on localities from the western Atlantic (Cape Cod (Cco), Western Bank (Web), Sable Bank (Sab), Trinity Bay (Tri), Southern Grand Banks (Sgb)) and the eastern Atlantic, deep sea frontal fish from the Barents Sea (BBBar), Iceland (BBIce), and Greenland (BBGre), intermediate fish from Iceland (ABIce) and Greenland (ABGre) and finally shallow water coastal fish from the White Sea (Whi), Iceland (AAIce), Faroe Islands (Far), Norway (Nor), North Sea (Nse), Celtic Sea (Cel), and the western and eastern Baltic (W-Bal and E-Bal).

**Supplemental Figure 6.**
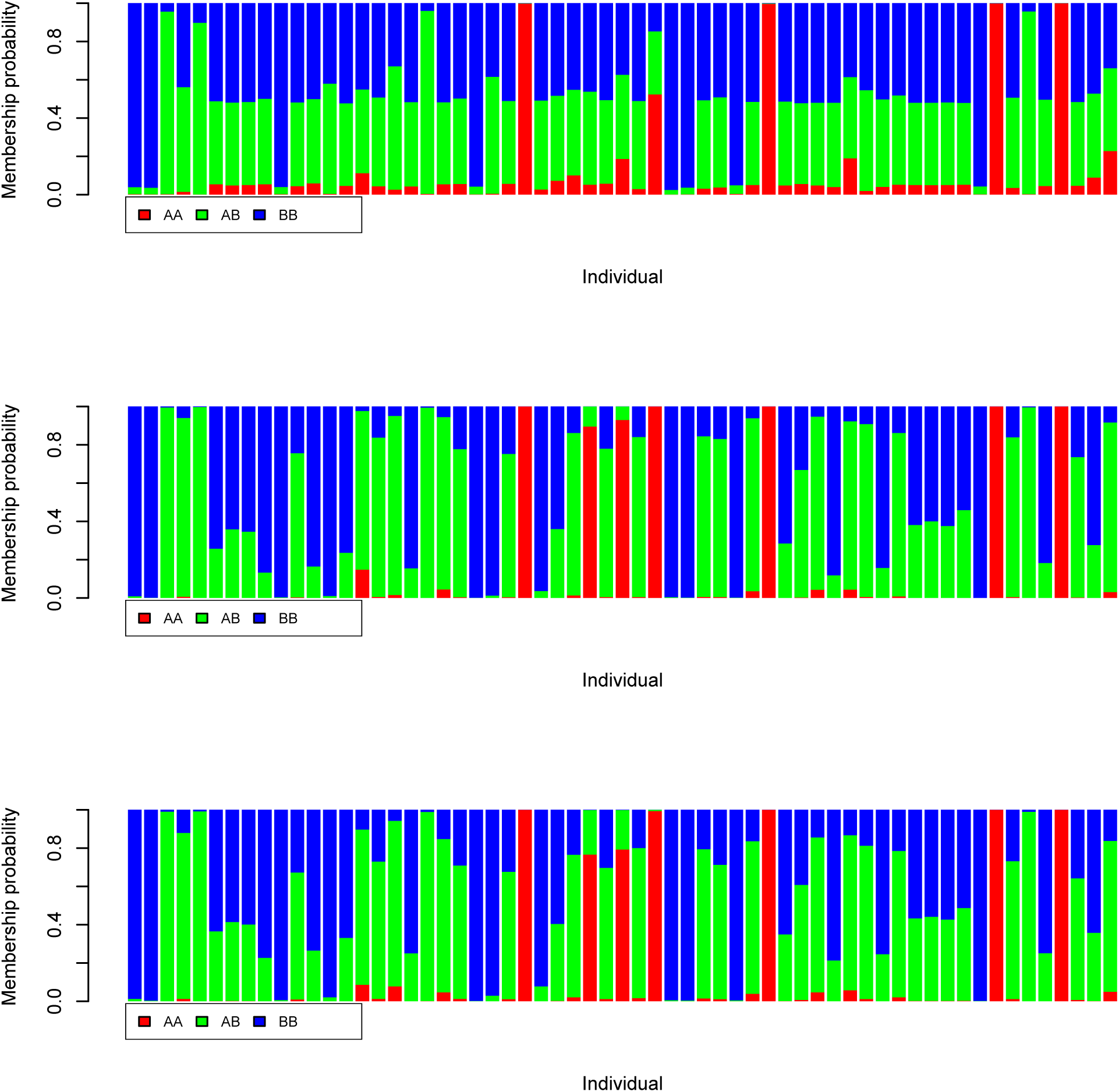
Posterior membership probabilities from the discriminant analysis of principal components (DAPC) in Figure 5. Genomic data that map to LG02 to LG23 (excluding LG01, top panel), data that map to all linkage groups LG01 to LG23 (middle panel), and data that do not map to linkage groups (bottom panel).

**Supplemental Figure 7.**
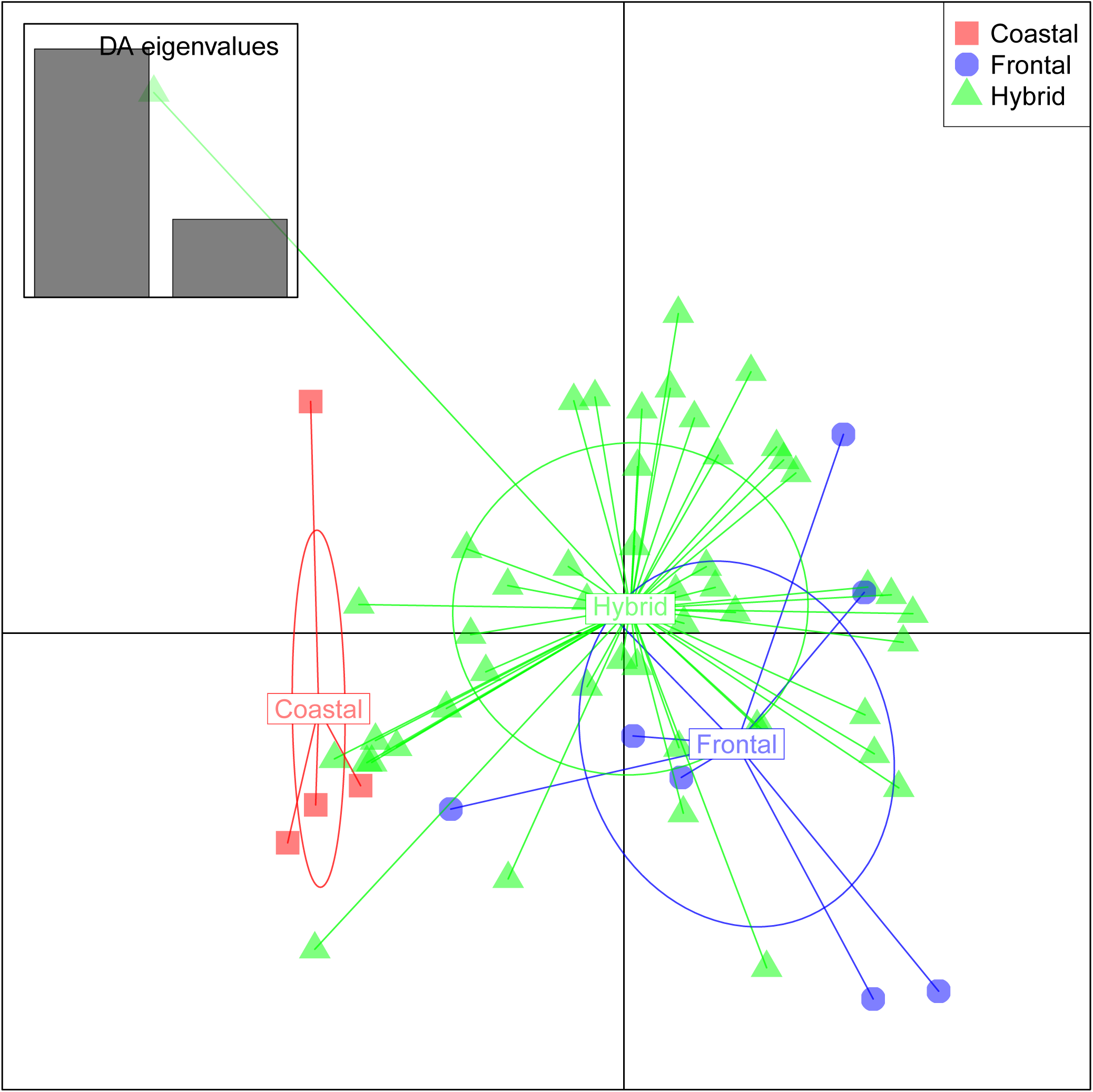
Discriminant analysis of principal components (DAPC) of phenotypic and habitat variation. Individuals were classified as pure coastal, pure frontal, and hybrid species by posterior membership probabilities of the DAPC analysis of linkage groups LG02 to LG23, the genomic parts that definately do not map to linkage group LG01 (Supplemental Figure 6 top panel). Phenotypic data are length, age, sex, ungutted and gutted weight, weight of liver, yearclass, and habitat data on depth.

